# Rapid divergence of male and female mitochondrial genomes in a basal protobranch bivalve *Yoldia hyperborea*

**DOI:** 10.1101/2025.06.07.658435

**Authors:** Dmitriy A. Fedorov, Alexandra V. Bezmenova, Margarita A. Ezhova, Glafira D. Kolbasova, Alexander B. Tzetlin, Tatiana V. Neretina, Dmitry A. Knorre

## Abstract

Doubly uniparental inheritance (DUI) of mitochondrial DNA (mtDNA) is a mode of mtDNA transmission whereby the male mitogenome is inherited predominately by males and the female mitogenome is inherited by females. DUI is unique to bivalve mollusks and has been identified in all major bivalve clades, including Protobranchia, the most basal bivalve subclade. Meanwhile, the mechanisms underlying the emergence and evolution of DUI remain poorly understood. Here, we focus on the early stages of divergence of male and female mitochondrial genomes by sequencing them in a basal Protobranchia species *Yoldia hyperborea* where they are characterized by low level of divergence, suggesting a recent origin of this system. Despite the relatively high relatedness of female and male genomes, we find that they have undergone rapid divergence both on the sequence and the structural level, including an origin of a novel gene, the ORFan, in the male genome. This novel gene is functional, as evidenced by the fact that it is transcribed and is subject to purifying selection. We show that this gene is a result of duplication and rapid evolution of another mitochondrial gene, NAD2. Additionally, the female mitogenome carries a duplicated cassette containing the cytochrome b gene, resulting in two different transcripts in female gonads both of which are subject to purifying selection, although their transcription levels differ by two orders of magnitude. These results showcase the rapid evolution of mitochondrial genes following their duplication, which occurred near the time of divergence of M- and F- mitochondrial genomes within a doubly uniparental inheritance system.

## Introduction

In most characterised metazoan species, mitochondrial genomes are inherited maternally (Birky 2001), whereas paternal mtDNA molecules are usually eliminated (Sato and Sato 2013; Sato et al. 2018; Lee et al. 2023). In rare cases, a small proportion of paternal mtDNA can be transmitted to the progeny (Kvist et al. 2003; Mastrantonio et al. 2019). However, some bivalve mollusks represent a striking exception to these rules, exhibiting an unusual pattern of mtDNA inheritance known as doubly uniparental inheritance (DUI). In the case of DUI, maternal (F-) mtDNA is transmitted to female germline and somatic cells, whereas the paternal (M-) mtDNA is restricted to male germline cells (Breton et al. 2007).

DUI bivalves are characterised by the presence of two divergent mitochondrial DNA variants (mitotypes) within populations and individuals. Two mitotypes are routinely found in DUI males (Fisher and Skibinski 1990; Hoeh et al. 1991), whereas adult female bivalves contain either only the F-type mtDNA or retain the M-type mtDNA at low levels (Sano et al. 2007). In DUI bivalves, the gametes—eggs and sperm—contain predominantly only one of the mtDNA variants (Skibinski et al. 1994; Venetis et al. 2006; Sano et al. 2007). Somatic cells of both males and females mainly contain the F-mtDNA; however, in some cases, both M- and F- mtDNA variants can be detected (Skibinski et al. 1994; Venetis et al. 2006). In a zygote, and during the early stages of embryonic development, paternal mitochondria remain aggregated. As a result, M-mtDNA is sequestered within a zone which supposedly develops into the male gonads (Cao et al. 2004). This segregation ensures transmission of M-mitochondrial DNA exclusively to male progeny (Zouros et al. 1994).

Nucleotide divergence of M- and F- genome of the same DUI species often exceeds 20%, indicating ancient origins of these systems (Mizi et al. 2005). In some species of Unionida, divergence of the M- and F- mitochondrial genomes predated species divergence (Rawson and Hilbish 1995). Moreover, some species have sex specific mtDNA regions which are present in the mitochondrial genome of one sex but not the other (Mizi et al. 2005). For example, the M-genomes of some species of the order Unionida contain an in-frame extension of the *COX2* gene of up to 200 codons (Curole and Kocher 2002); furthermore, *Scrobicularia plana* (order Cardiida) encodes a Cox2 protein with a C-terminal extension of 1892 a.a. (Capt et al. 2020; Tassé et al. 2022). Mitochondrial genomes of some bivalve species contain sex specific ORFans, putative protein coding regions without clear homology to any other characterised proteins (Milani et al. 2013).

DUI enables long-term coexistence of two different mitochondrial genomes in germline which creates unique conditions for their evolution. First, the presence of two genomes weakens purifying selection for one of them, the M-type mitogenome. The M-type genomes have higher substitution rates and accumulate substantially more non-synonymous amino acid substitutions than the F-genomes (Hoeh et al. 2002; Ort and Pogson 2007). Furthermore, the M-type mitogenome of *Donax* species lacks several core protein coding genes (PCGs) encoding subunits of respiratory complex I (Burzyński et al. 2024), exemplifying relaxation of selection pressure on the M-type mitogenomes. Second, the DUI system suggests co-occurrence of M-type and F-type mitogenomes in the same germline cells, for example in zygote. This makes mtDNA recombination possible; indeed, multiple recombination events between M- and F- type mitogenomes were reported in mussels (Ladoukakis and Zouros 2001).

DUI has been found in over 100 species (Gusman et al. 2016). Importantly, based on standard mitochondrial marker genes, two studies reported DUI in *Yoldia hyperborea* and *Ledella spp* (Boyle and Etter 2013; Gusman et al. 2016). *Yoldia hyperborea* and *Ledella spp* species belong to Protobranchia, a basal subclass of bivalves (Crouch et al. 2021). However, many other bivalves show low divergence between mtDNAs isolated from male and female gonads, suggesting that one of the genomes could displace the other in some clades (Doucet-Beaupré et al. 2010).

In this study, we describe a case of recent divergence of M- and F- type mitogenomes in Protobranchia, a basal group of bivalves. For this, we sequenced and assembled mitochondrial genomes from male and female gonads of two Nuculanida (Protobranchia) species, *Yoldia hyperborea* and *Nucullana pernula*. While we found only minor differences between mitochondrial genomes of male and female *Nucullana pernula* specimens, the male and female mitogenomes of *Yoldia hyperborea* exhibited pronounced divergence despite their young age. In particular, male and female mitogenomes are characterised by the presence of pseudogenes and genes that are absent in the complementary mitogenome as well as an elevated rate of nonsynonymous evolution, indicating rapid functional divergence of the two.

## Results

### Structural variation in Protobranchia mitogenomes

To determine the differences between male and female mitochondrial genomes of protobranchia bivalves, we collected four specimens: male and female individuals of *Yoldia hyperborea* and *Nucullana pernula,* and isolated DNA from their gonads (see Material and Methods). Combined Illumina short-read and PromethION long-read sequencing of their total DNA enabled us to assemble four corresponding mitochondrial genomes. All genomes contained the core set of mitochondrial genes typical for metazoan mitochondrial genomes: 13 PCGs, 2 rRNAs, and 21 tRNAs (22 in F-mitogenome of *Y. hyperborea*). All PCGs were encoded on the same strand. Consequently, the DNA sequence displayed a pronounced strand asymmetry, characterised by a positive GC-skew and a negative AT-skew (Figure 1).

**Figure 1.**
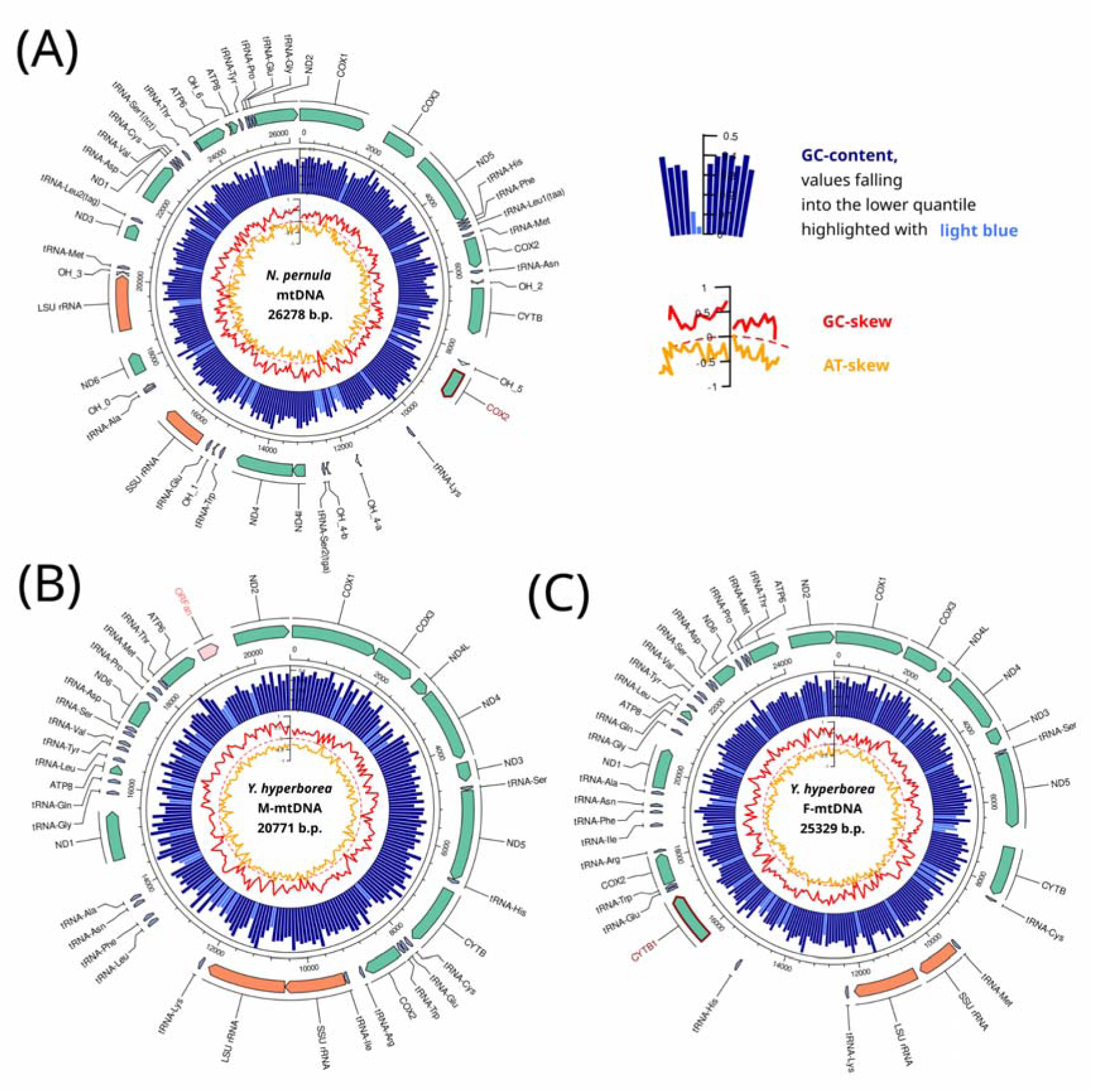
Annotated map of mitochondrial *Nuculana pernula* (A) and *Yoldia hyperborea* (B, C) genomes. The outer track represents the boundaries of standard mitochondrial protein coding genes, the ORFan, tRNAs and rRNAs. Direction of the arrow corresponds to the coding strand. Blue histogram shows the GC-content, with light blue highlighting the 10% most AT-rich regions. Red and yellow, nucleotide GC- and AT-skew correspondingly.

The mitochondrial genomes obtained from the female and the male individuals of *N. pernula* were very similar (see below). Therefore, we pooled reads from them to obtain a single mitochondrial genome, which was 26,278 bp in length. Notable features of the *N. pernula* genome were the separation of SSU and LSU rRNA genes by tRNA-Ala and the *ND6* gene and the duplication of the *COX2* gene (Figure 1A). One *COX2* paralog is located at 5296:5980 bp, while the second paralog has coordinates 8503:9180 bp. Despite the presence of two copies of the gene, the second paralog has a stop-codon inside the sequence which breaks it into two parts.

Unlike *N. pernula,* assemblies from male and female *Y. hyperborea* yielded highly different genomes, respectively referred to as M- and F- genomes. The differences were observed both in genome size (M-genome: 20,771 bp; F-genome: 25,329 bp) and architecture (Figure 1B,C). Architectures of both M- and F- genomes differed from the *N. pernula* mitogenome and from each other, as detailed below.

First, the F-genome of *Y. hyperborea* contained two clearly identifiable *CYTB* sequences, while only one *CYTB* was identified in the M-genome (Figure 1C), indicating a duplication of *CYTB* In the female genome. One of the paralogs *(CYTB)* has coordinates 7062:8195 bp, while the other (hereafter, *CYTB1)* is located in the region 15916:17064 bp (Figure 1C). Both paralogs of the gene have similar lengths, but differ in nucleotide composition. The second paralog is located next to the *COX2* gene. To time *CYTB* duplication, we aligned the *CYTB* genes and reconstructed a phylogenetic tree (Figure 2). The phylogeny reconstructed using synonymous sites clustered *CYTB* and *CYTB1* from the F-mitogenome in a single clade (Figure 2B), while the phylogeny based on non-synonymous sites placed the duplicated *CYTB1* gene at the root of the F- and M-*CYTB* genes (Figure 2C). This indicates that *CYTB1* duplication occurred either shortly before or shortly after the divergence of the *Y. hyperborea* F- and M-mitogenomes.

**Figure 2.**
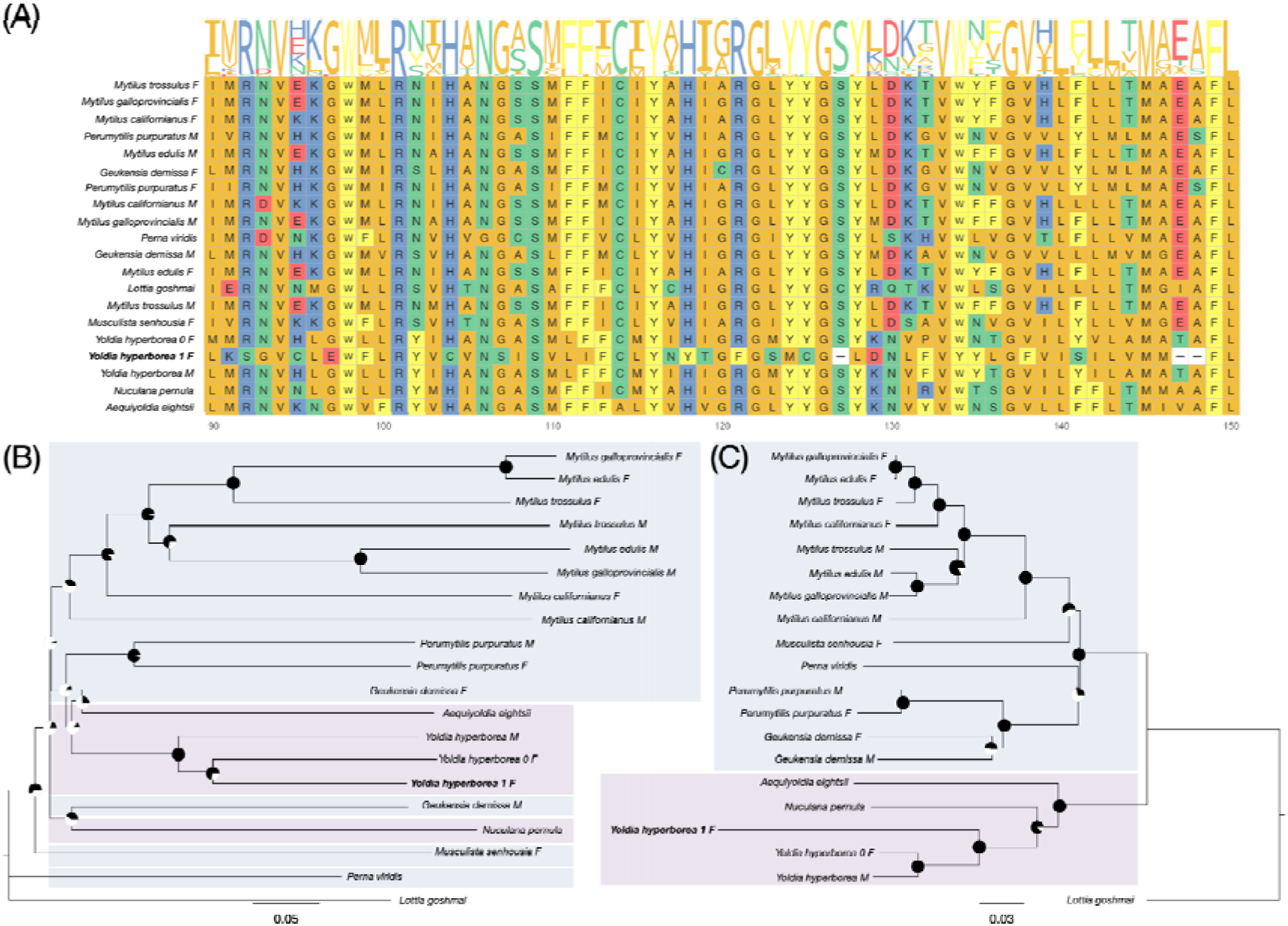
Duplication of *CYTB* gene in the F-genome of *Y. hyperborea*. (A) Multiple alignment of a region of *CYTB* proteins showing divergent sequence of *CYTB1* gene in the female mitochondrial genome of *Y. hyperborea*. Note that the highlighted sequence has nucleotide deletions in multiples of three. (B, C) The phylogenetic tree of *CYTB* gene sequences constructed using the NJ algorithm with 100 bootstrap replicates(indicated as black area on the node circles), based on synonymous (B) or nonsynonymous (C) substitutions only. Numbers of differences per site were calculated using the Nei-Gojobori method. Note that for synonymous substitutions (B), both genes of *Y. hyperborea* represent a monophyletic clade with similar branch lengths for each gene, while for nonsynonymous substitutions (C), the branch length for *CYTB1* gene from *Y. hyperborea* is greater than that for any other *CYTB1* gene in the sample.

Second, the M-genome of *Y. hyperborea* harboured a putative protein-coding region of 363 bp (coordinates 19028:19391 bp) downstream of the *ATP6* gene, lacking stop codons in one reading frame of 121 amino acids which we refer to as the ORFan (Figure 1A). While the length of ORFan is approximately half of that of other *NAD2* genes, its alignment with *NAD2* genes of *Y. hyperborea* as well as other bivalves contains extended conservative regions (Figure S1, Table S1). A weak homology between ORFan and *NAD2* indicates common ancestry between them.

Third, the genomic position of the *COX2* gene differs between the female and male genomes of *Y. hyperborea.* In the M-genome of *Y. hyperborea*, the *COX2* gene has coordinates 8245:8940 bp, while in the F-genome it is located in the region 17416:18111 bp. In the region in which the *COX2* gene is located in the male, there is a region in the female without detectable homology based on the genome alignment (Figure 3A). However, if we separately align the *COX2* gene in the male against this region in the female, we find an unexpectedly high number of coincident nucleotides, despite the presence of numerous insertions and deletions not in multiples of three. This suggests that the similarity of these regions is not accidental. To test this, we simulated sequences of the same GC or codon content as that of the considered region in the female by either (i) generating a random sequence with the same nucleotide content as that of the male *COX2* gene, or (ii) reshuffling the codons of the male *COX2* gene, and then aligning the resulting artificial sequences to the considered region of the female genome. In all cases, when comparing the percentage of identical positions in the real and artificial sequences, the alignment with the real sequence produced much higher nucleotide similarity in the alignment (Figure 3B). This indicates that the region of the F-mitogenome syntenic to the male *COX2* locus contains a highly degraded *COX2* pseudogene with numerous short indels of random lengths.

**Figure 3.**
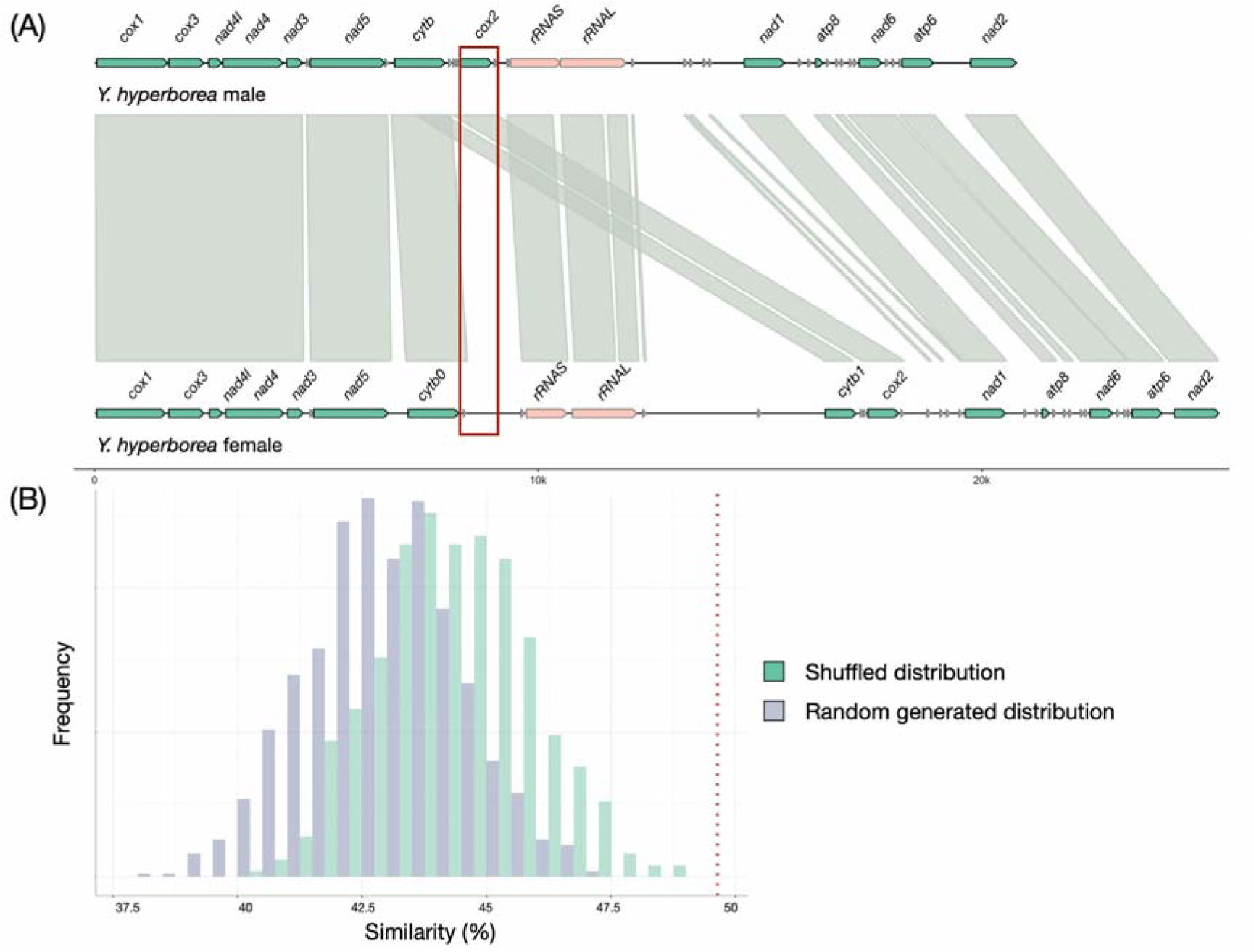
Remnants of *cox2*. (A) Genome alignment of male and female *Y. hyperborea* genomes with homology blocks shown by olive bars. Mitochondrial genes are shown with arrows, with the arrow color corresponding to gene class. The red box indicates a hypothetical site of *COX2* pseudogenization. (B) Histogram of % sequence similarity between the shuffled (green) or randomly generated (gray) sequences of the male *COX2* gene and the colocalized region of the female genome. The percent of similarity between *COX2* gene and original sequence is shown with a red dashed line.

### Both M- and F- genes are expressed in male and female Yoldia

To study gene expression, we isolated RNA from the gonads of male and female *Y. hyperborea* and *N. pernula*. We then obtained cDNA using polyT primers for reverse transcription, prepared libraries, and sequenced them using the Illumina platform (see Materials and Methods). The resulting transcriptome reads were mapped to the genomic assemblies, and we estimated the coverage of each mitochondrial gene in males and females. Only high-quality reads that clearly mapped to the assemblies were selected for analysis. Notably, the level of divergence between M- and F- type genomes was sufficient for a clear distinction between the M- and F- type RNAseq reads (Figure 4A). As expected, we found that the F-reads were predominant in the F-genome, and the M-reads were predominant in the M-genome (Figure 4B).

**Figure 4.**
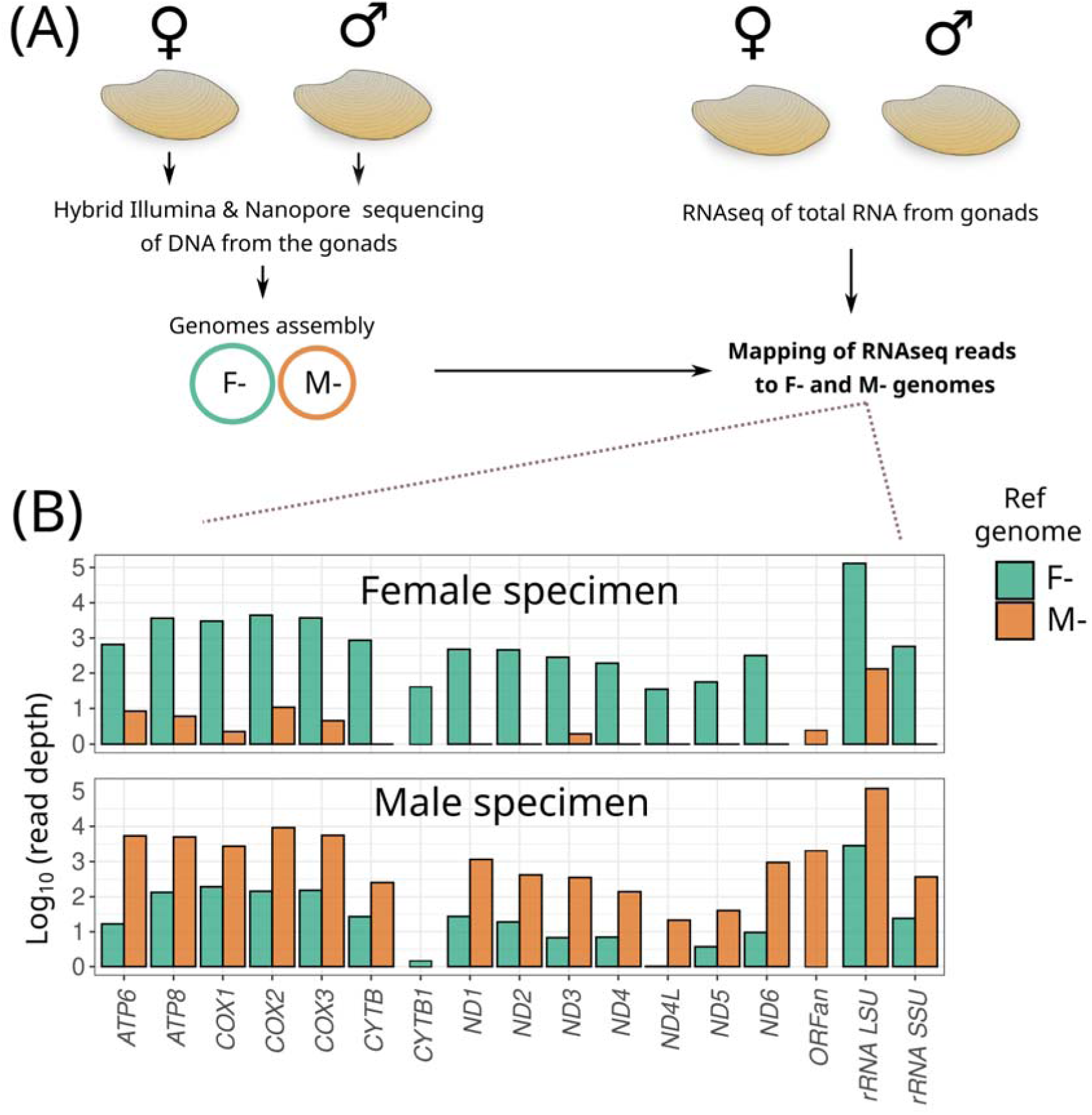
Relative expression of mitochondrial genes, ORFans and NCRs in mitochondrial genomes of male and female specimens of *Yoldia hyperborea*. (A) Schematic workflow for mapping RNA-seq reads to M and F genomes. (B) Expression of M and F genes in male and female gonads, shown as log10(read depth per base pair). For comparison, expression levels in female gonads were normalised by dividing all values by 0.9006 to account for the smaller total number of reads in this preparation.

Transcriptomic data confirmed transcription of both duplicated genes, ORFan and CYTB1. In fact, the expression level of the ORFan in the male gonads exceeded the expression of several standard PCGs, e.g. *NAD3* and *NAD4L* (Figure 4B). Importantly, the ORFan transcript was not a part of the upstream PCG transcript, as evident by the drop in coverage between the *ATP6* gene and the ORFan (Figure S2). Similarly, we identified the transcripts of the F-genome duplicated *CYTB1* gene. Its expression level was an order of magnitude lower than that of the original *CYTB* (Figure 4B, Figure S3); nonetheless, in the female genome, it was expressed at a level similar to some of the *NAD* gene transcripts. Taken together, these observations show that the duplicated genes, *CYTB1* and ORFan, produce mRNA that is present in quantities similar to some other mitochondria PCGs.

Consistent with previous findings (Skibinski et al. 1994; Venetis et al. 2006; Sano et al. 2007), we observed significantly higher expression of male mitochondrial genes in males and female mitochondrial genes in females. Still, female variants of mitochondrial genes, as well as the female-specific *CYTB1* gene, were also expressed in males, albeit at much lower levels than the male variants. Notably, we also detected expression of male mitochondrial gene variants, as well as the male-specific ORFan, in females (Figures 4 and 5). While these expression levels are generally low for most genes, they are supported by tens to hundreds of reads.

**Figure 5.**
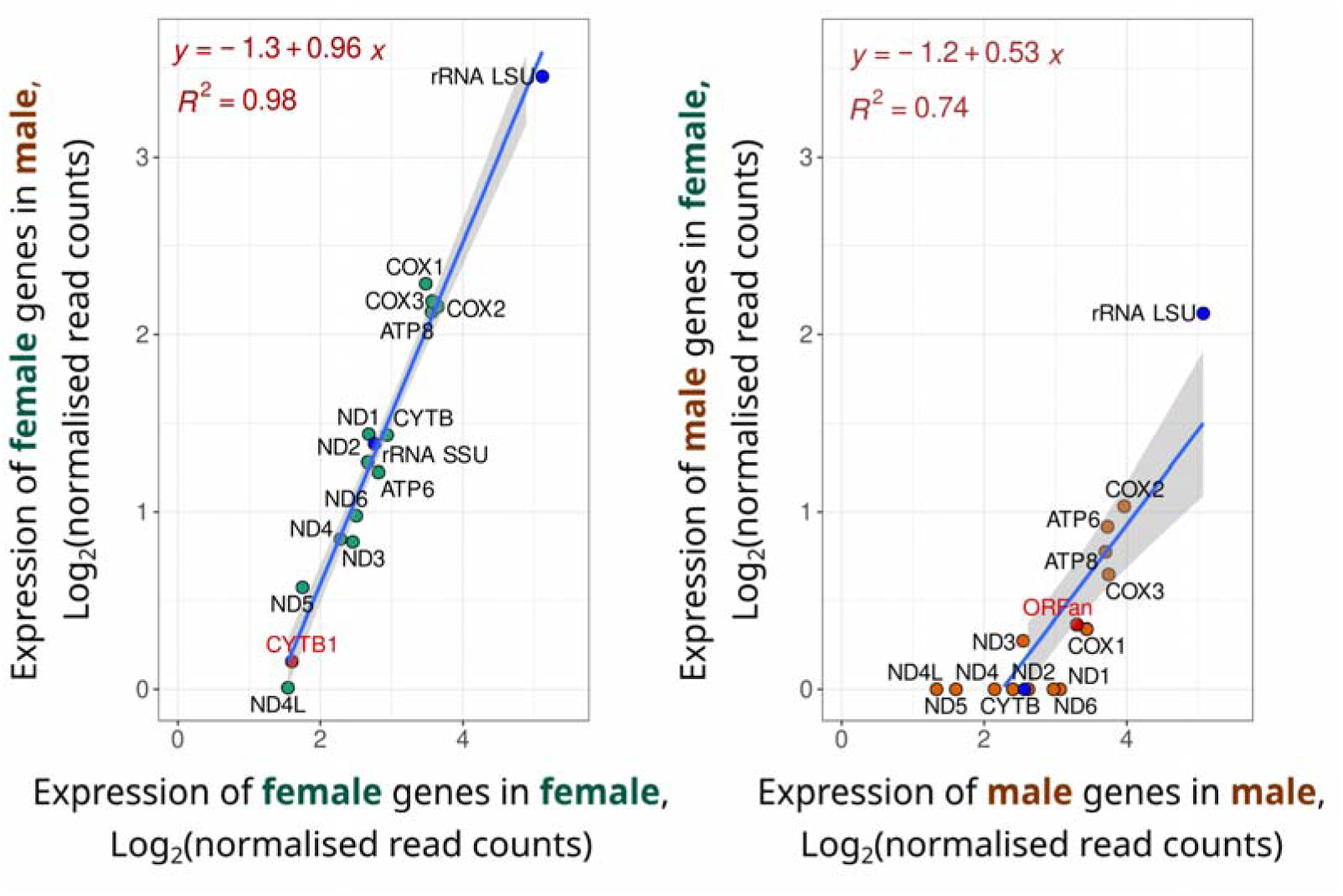
The mitochondrial gene expression level in the organism of the opposite sex is maintained in a similar proportion relative to the main sex.

It has been hypothesized that the expression level of some of the mitochondrial genes may play a role in sex determination in a DUI system (Breton et al. 2011). In particular, this role could be played by the sex-specific mitochondrial genes, *CYTB1* or ORFan. If so, we could expect the relative expression of individual genes, such as ORFan, to differ between male and female mitochondria. Therefore, we hypothesized that the expression levels of these genes relative to that of other genes would differ between RNA preparations isolated from males and females. To test this, we constructed linear regressions comparing the expression levels of M genes in male and female gonads, and similarly for F genes. As shown in Figure 5, there is a nearly perfect correlation between the expression levels of F genes, including *CYTB1,* in male and female gonads. The expression of M genes, including ORFan, also shows a strong linear relationship between male and female genomes. This result argues against sex-specific regulation of expression of mitochondrial genes.

### Sequence divergence in M- and F- genomes of Y. hyperborea

The presence of two types of mtDNA transcripts in each species enabled us to compare relative distances between the two M- and the two F-genomes in *Y. hyperborea.* For this, we filtered the RNAseq reads mapped to the highly expressed regions of the mitogenome (see Material and methods and trans_distance_male_female tab in Dataset1.xls). The divergence between the M-genome in males and females, as well as the F-genome in males and females, was minor (Figure 6), probably reflecting intrapopulation variation. By contrast, the nucleotide sequence divergence between the *Y. hyperborea* male and female mitochondrial genomes was much higher, and also by far exceeded that in *N. pernula,* indicating DUI in *Y. hyperborea* but not in *N. pernula* (Figure 6).

**Figure 6.**
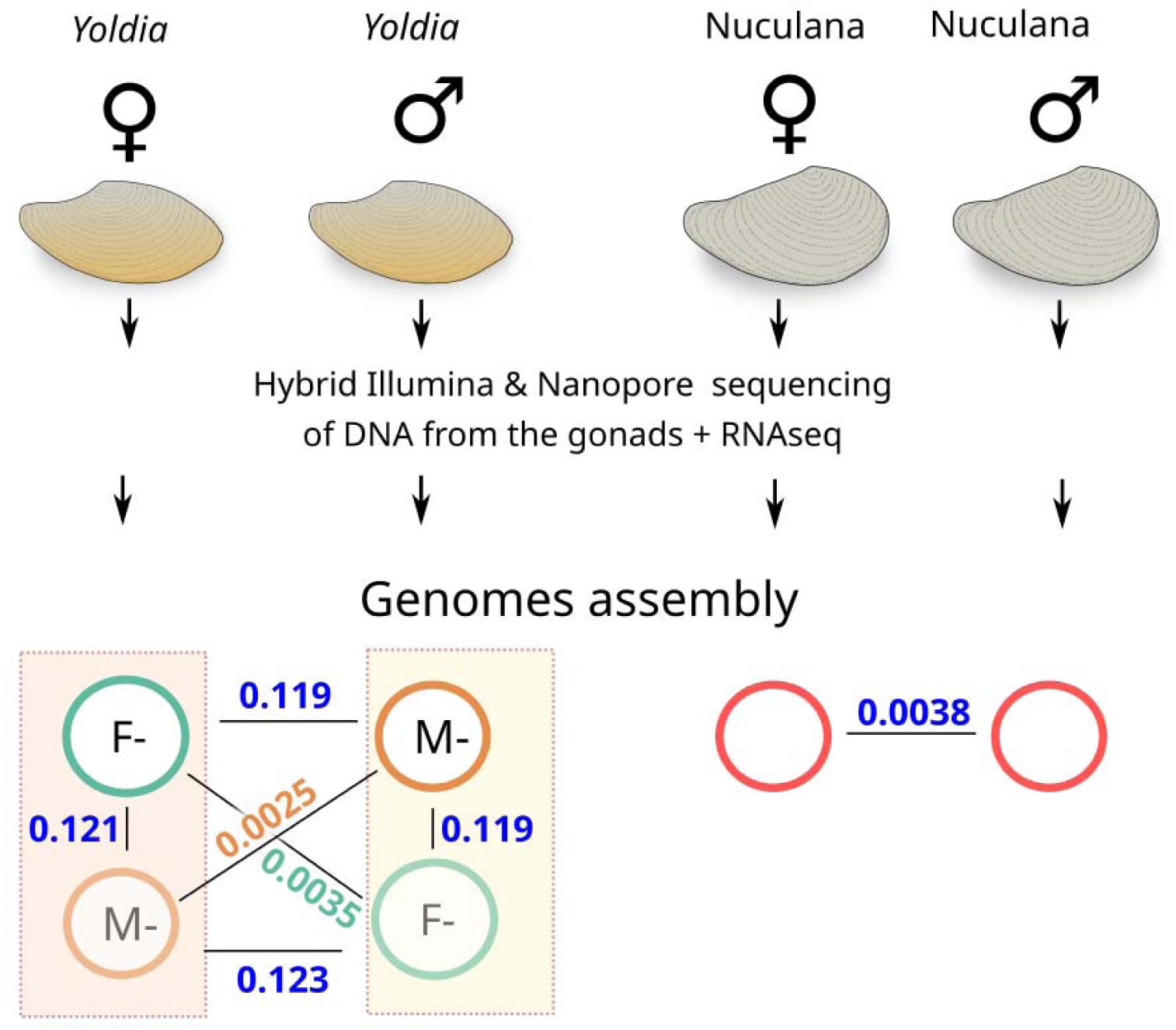
Relatedness of assembled Protobranchia mitogenomes. Relative frequency reflects the proportion of reads mapped to F- and M-mitogenomes from DNA samples isolated from male and female bivalves. Divergence was calculated as mean similarity between several mitochondrial genes (*cox1, cox2, cox3, rRNAL*) identified in male and female transcriptomes. For minor mitochondrial variants (i.e. male mitochondrion in female transcriptome and female mitochondrion in male transcriptome) reads strictly mapped to such variants were mapped on the previously assembled genome and consensus sequence with snips was further generated.

To evaluate the level of divergence between the M- and F- genomes of *Y. hyperborea,* we measured nucleotide (Figures 7) and amino-acid (Figure S4) distances for all PCGs and compared them with those for other DUI bivalves with verified complete mitochondrial genomes in GenBank. We excluded the *ATP8* gene from analysis because of its shortness and low conservation which prevent robust alignment. *Y. hyperborea* had the lowest nucleotide sequence distances between the M- and F- genomes among the selected mitogenomes, indicating that this is the DUI system with a low divergence (Figure 7). The presence of distinct male and female mitogenomes in *Y. hyperborea* but not in the closely related *N. pernula,* together with the low evolutionary distances between the *Yoldia* male and female mitogenomes, indicate a recent divergence of the DUI system in this clade of Protobranchia.

**Figure 7.**
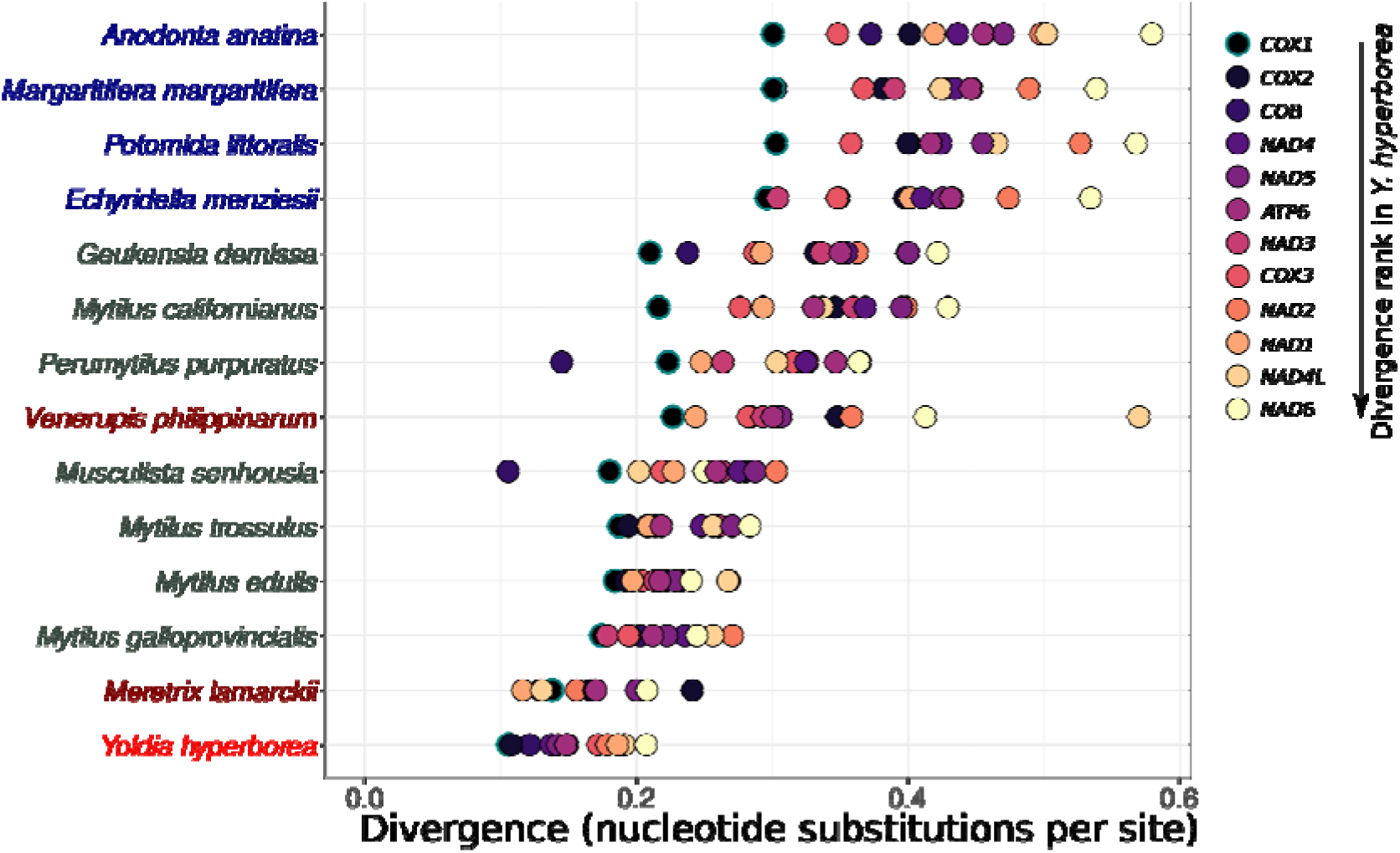
Mitochondrial protein-coding gene (PCG) sequence divergence between M- and F- mitogenomes of DUI bivalves. Each data point represents the nucleotide divergence of an individual PCG between Mand F-genomes of the same species. Species are ordered by average nucleotide distances of PCGs and colour-coded according to their taxa

### Relaxed selection on amino acid sequence in the male genome of *Y. hyperborea*

We reconstructed a phylogenetic tree from the concatenated protein-coding genes of bivalve species from clades at a range of phylogenetic distances (Figure 8). The tree confirmed the basal position of Protobranchia bivalves, including *Yoldia* and *Nuculana* as well as *Aequiyoldia eightsii*, the protein coding sequences of which were assembled from available RNAseq results (SAMN32616087). Notably, the M- and F- mitogenomes of *Yoldia* clustered together, confirming a relatively recent divergence. This contrasts with Unionids, where the divergence of M- and F- genomes preceded speciation.

**Figure 8.**
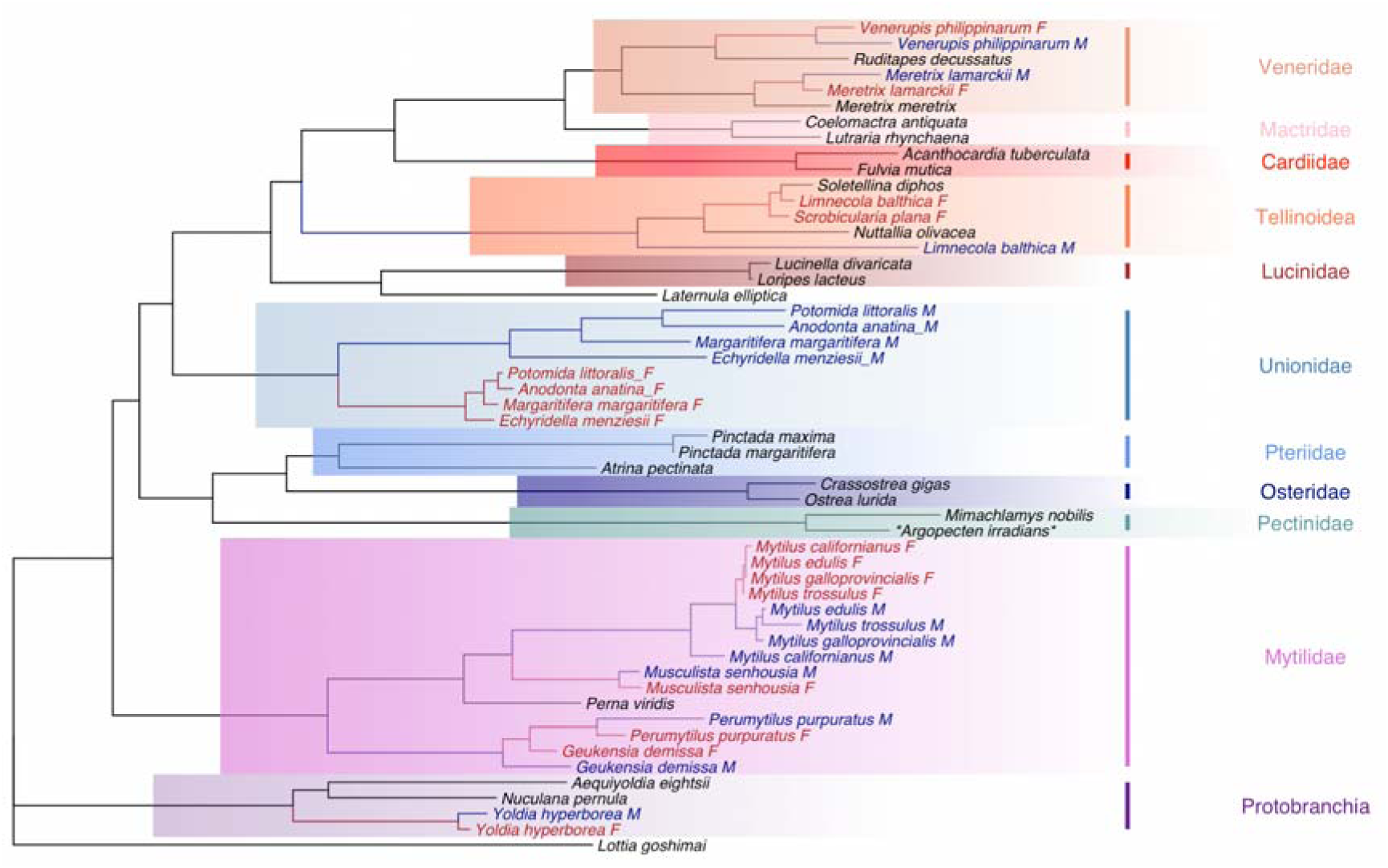
Phylogenetic tree of bivalves with known sex-specific M- and F- genomes. Phylogenetic tree of 53 bivalve mitogenomes based on coding sequences of 12 standard protein-coding genes, excluding ATP8 (concatenated codon multiple sequence alignments of PCGs). The tree was constructed using the maximum likelihood (ML) method with the mtZOA+R4 substitution model. Numbers represent bootstrap values determined by 1000 ultrafast bootstrap replicates. The tree is rooted with the *Lottia goshimai* mitogenome as an outgroup. For those species whose names are not followed by the letter M or F, the DUI is not described. The detailed tree with bootstrap support and branch length is presented in File S2.

To investigate the intensity of selection on the M- and F- genomes, we calculated the Omega (dN/dS) value across the concatenated set of protein coding mitochondrial genes over the evolutionary lineages leading to each such genome. To obtain the Omega values most characteristic of the mitogenome-specific evolution, we only calculated it over the substitutions in the terminal branches of the phylogeny. Omega was clearly elevated for the male mitogenomes, compared to the female mitogenomes (Figure 9). In every single pair of M- and F- mitogenomes of the same species, the dN/dS value was higher in the M- mitogenome. Furthermore, we observe a negative association between the Omega value for the substitutions discriminating between the male and female mitochondrial genomes and the phylogenetic distance between them (Figure S5).

**Figure 9.**
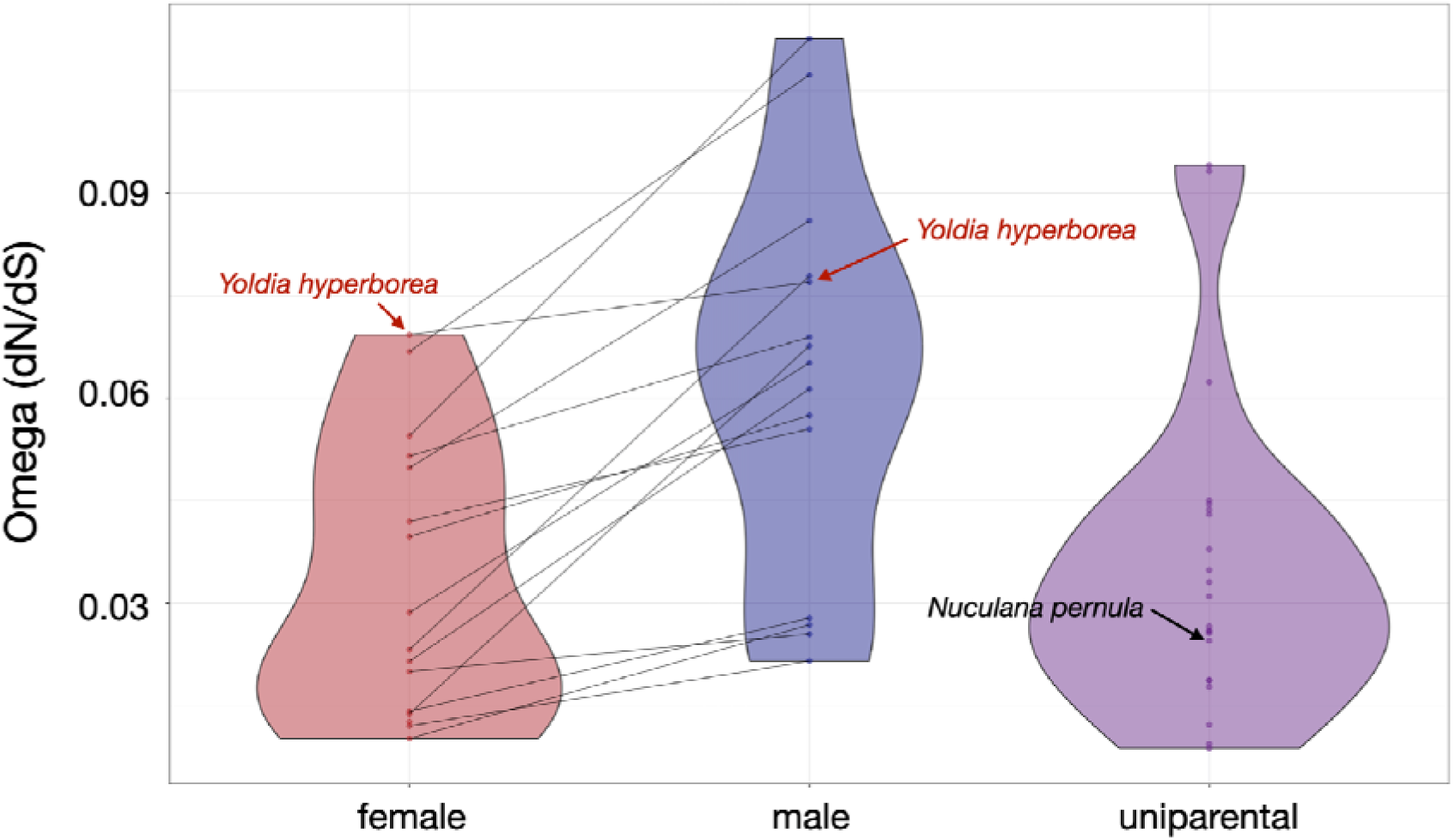
dN/dS values in mitochondrial genes across various bivalve species. Omega (dN/dS) was estimated for a concatenated set of all mitochondrial protein-coding genes (excluding *ATP8*) with a free-branch model using codeml from paml package. Lines connect values for the same species but different sex types (F- or M- respectively). The dN/dS values for male genes are elevated for every single pair of studied genomes (Figure 8).

To estimate the overall difference in selection between the M- and F- mitogenomes, we concatenated alignments of protein coding genes and compared the fit of two types of models: a single Omega value (codeml model 0) and separate Omega values for the three types of branches: those leading to the M-mitogenomes, those leading to the F- mitogenomes, and the remaining branches (i.e., those leading to uniparental mitogenomes or internal branches; codeml model 2). The model allowing for different Omega values (model 2) had a much better fit (Table S2), indicating a strong difference in Omega values between branch types. Indeed, the Omega of the M-branches was ∼2.2x that of the F-branches (0.085 vs. 0.039). Together, these results indicate strongly relaxed negative and/or positive selection in the M-mitogenome, compared to the F-mitogenome. These results also largely held for alignments of individual genes (Table S3).

The female genome contains two paralogs of the *CYTB* gene (Figure 1С), and the difference in their branch lengths between synonymous and nonsynonymous sites (Figure 2) indicates a change in selection between them. To study this, we measured the Omega values for the two paralogs in *Y. hyperborea* individually, and compared them to those of other species. The Omega of the CYTB gene in *Y. hyperborea* was similar to that for this gene in the rest of the tree. By contrast, the Omega of the CYTB1 was ∼4x higher than that of the CYTB genes in other species (Table 1), indicating relaxed negative selection or positive selection acting on CYTB1 in the F-genome.

**Table 1.** The omega (dN/dS) values estimated for the two paralogs of CYTB in the female *Y. hyperborea* genome.

|  |  |  |  |  |  |  |
| --- | --- | --- | --- | --- | --- | --- |
| Gene | w0<br>model | w2<br>(CYTB, *Y.*<br>*hyperbor*<br>*ea*) | w2 (rest) | -lnL (w0) | -lnL (w2) | P-value |
| *CYTB* | 0.04202 | 0.04166 | 0.0492383 | -31417.837728 | -<br>31417.75083<br>1 | 0.677 |
| *CYTB1* | 0.04250 | 0.15872 | 0.03866 | -31068.617458 | -<br>31047.99968<br>3 | 1.35<br>$\cdot 10^{-10}$ |

For each gene, we compare the fit of the model assuming a single Omega value across the phylogenetic tree (w0) to a model allowing for an Omega value specific to the considered gene (*CYTB* or *CYTB1*) of *Y. hyperborea* F-mitogenome (w2). P-value corresponds to the significance in the differences between the two models.

## Discussion

Many species of bivalve mollusks have divergent gender-linked mitochondrial genomes (Gusman et al. 2016). This high level of mtDNA divergence (see Figure 8) indicates the presence of doubly uniparental inheritance (DUI) of mtDNA and rules out recent recombinations between M- and F- genomes. Conversely, male and female specimens of some other bivalves possess more similar mitogenomes, suggesting either strict uniparental inheritance (UI) of mtDNA or a recent event of masculinization of F-mtDNA within a DUI context. The masculinization of F-mtDNA implies that M-mtDNA has been replaced by F-mtDNA in DUI species, making it indistinguishable from species with UI of mtDNA (Hoeh et al. 1997). Although the feminization of M-mtDNA in certain clades cannot be ruled out, it is considered highly unlikely based on phylogenetic reconstructions of M- and F- genomes (Hoeh et al. 1997). Additionally, feminization of M-mtDNA is unlikely because M-mitochondrial genomes are typically much less fit than maternally inherited mitochondrial DNA. Indeed, individuals of DUI species *Arctica islandica* with M- type mtDNA in somatic cells exhibit decreased mitochondrial activity (Dégletagne et al. 2021), and spermatozoa of DUI species with M- type mtDNA are less motile than those of UI species (Bettinazzi et al. 2020).

Here we examined two related bivalve species with high (*Y. hyperborea*) and low (*N. pernula*) divergence of mitochondrial genomes (Figure 1). These species belong to Protobranchia, a basal group of bivalves (Figure 8). In line with previous studies (Hoeh et al. 1996; Doucet-Beaupré et al. 2010; Gusman et al. 2016), our phylogenetic reconstruction does not result in distinct clusters grouping the M- or F-mitogenomes of different species together, similar to those observed for *Unionidae* or *Mytilus sp.* (Figure 8). This is consistent with two scenarios: (i) multiple independent origins of DUI in different clades, and (ii) a monophyletic origin of DUI in an ancestral bivalve with repeated losses in multiple clades. The second scenario could occur either through a change in the inheritance pattern or through masculinization of the F-genome. Although our study cannot definitively distinguish between these possibilities, the high prevalence of DUI among the bivalves, including *Protobranchia*, supports the idea that DUI is either ancestral to all bivalves or that all bivalves have a preadaptation enabling the transition from UI to DUI.

Our study revealed several unusual features of the mitochondrial genomes in Protobranchia bivalves. First, we identified the presence of M-genome transcripts in female gonad samples (Figure 4). While the presence of the F-genome in male somatic cells is a common characteristic of DUI species (Fisher and Skibinski 1990; Hoeh et al. 1991), the reverse situation—the presence of the M-genome in female somatic cells—is less common (Sano et al. 2007). Importantly, our preparations from gonads may contain germline and somatic cells DNA. The presence of M-mtDNA in somatic cells of DUI species has also been reported in the female somatic tissues of *Arctica islandica* (Dégletagne et al. 2021). Similar to *Yoldia*, this species is characterized by one of the lowest levels of divergence between F- and M-mitochondrial genomes, exhibiting only a 5.5% difference in DNA sequence (Dégletagne et al. 2016; Dégletagne et al. 2021). The presence of M-genomes in female specimens of *Y.hyperborea* and *A. islandica* suggests that the sorting efficiency of M- and F- genomes might be complicated in species with low divergence between these genomes. We suggest that this may contribute to the increased frequency of F-genome masculinization in species with low M- and F- genome divergence.

Second, we did not detect divergent mitochondrial genomes in the gonads of *N. pernula*. While the absence of DUI is a common characteristic of bivalves, it is typically associated with hermaphroditism (Guerra et al. 2017). In contrast, *N. pernula* appears to be a rare example of a gonochoric species without apparent DUI (Figures 6 and 8), while DUI is uncommon among hermaphrodites. Exceptions to both rules have previously been identified, e.g. a gonochoric non-DUI unionid species *Mutela dubia (Guerra et al. 2017)*, and the DUI observed in the hermaphrodite *Semimytilus algosus* (Mytilida) (Lubośny et al. 2020). Collectively, the presence of non-DUI gonochoric species challenges the idea that sex determination in gonochoric bivalves is driven by mitochondrially encoded factors, although this does not entirely exclude this possibility in lower-level taxa such as fresh-water mussels where the DUI is strictly maintained (Breton et al. 2011).

Another distinctive feature of the *Y. hyperborea* mitogenomes is a high dN/dS values for both M- and F- genomes despite their recent divergence compared to other DUI bivalves (Figure 9). The low level of divergence between the male and female mitochondrial genomes of *Y. hyperborea* may indicate that they diverged recently, compared with known examples among freshwater bivalves (Unionidae), where the divergence occurred at the root of the group (Figure 8). Although the first stages of mitochondrial genome divergence within a single species remain hidden, at least in zygotes, two different genome variants should coexist in a single cell. Therefore, patterns of these genomes may be similar to the evolution of duplicated genomes or genes.

Accordingly, comparison of dN/dS of the bivalve species with DUI reveals a strong negative correlation between the proportion of nonsynonymous and synonymous substitutions fixed between the male and female mitochondrial genomes and the phylogenetic distance between them (Figure S5). In line with these results, a recent study of Veneridae bivalves with DUI found a similar pattern: the species with the lowest levels of the divergence exhibited highest dN/dS ratio (Xu et al. 2024). The negative correlation between dN/dS and divergence has been described previously in a range of systems, both in comparisons of orthologs and paralogs. It can originate from multiple causes, including technical artefacts (such as an excess of sequencing, mapping and/or alignment errors at low phylogenetic distances spuriously inflating dN), correlated mutation events, an excess of polymorphic (unfixed) nonsynonymous mutations at low phylogenetic distances, and an episode of relaxed negative or positive selection at early stages of speciation (Lynch and Conery 2000; Rocha et al. 2006; Kryazhimskiy and Plotkin 2008; Wolf et al. 2009; Stoletzki and Eyre-Walker 2011; Mugal et al. 2014; Stolyarova et al. 2019; Burskaia et al. 2020).

Irrespective of its cause, this pattern suggests that with increased divergence, the negative selection shaping the mitochondrial subgenomes asymptotically reaches a low value (dN/dS ∼ 0.01 to 0.1), underlying the strong selection on both the male and the female subgenomes.

Divergence of M- and F- mitogenomes is evident not only at the level of sequence but also in their structural features which evolve by duplication and rearrangements of genes and gene clusters. In our study, we detect three duplicated regions within M- and F- genomes of *Y. hyperborea* (Figure 10). First, we observed a duplication of the *CYTB* gene in the female mitochondrial genome (Figure 2). The duplicated copy of the *CYTB* gene copy contains an increased number of nonsynonymous mutations, which are subject to selection, while the number of synonymous mutations in the copies is equal (Figure 2), indicating relaxed negative and/or positive selection acting on it. Second, the weak homology of the *ORFan* in the male mitochondrial genome with the *NAD2* gene (Figure S1), as well as numerous frameshift and non-synonymous mutations in the *ORFan* region, may indicate that the *ORFan* is a highly diverged and partially degraded copy of the *NAD2* gene, which was formed after duplication. The question of whether the *ORFan* is translated into a protein remains unclear, but the presence of RNA-seq reads in the ORFan regions (Figure 1) suggests that the corresponding mtDNA region is transcribed and processed. Finally, the region that is apparently duplicated in the female mitochondrial genome of *Y. hyperborea* contains not only the *CYTB* gene, but also the *COX2* gene. The syntenic position of the *COX2* gene in the mitochondrial genome is not annotated as a gene in the mitochondrial genome (Figure 3). However, when aligning the male and female genomes, it can be seen that the percentage of similarity of this region with the syntenic region in the male mitochondrial genome, where the *COX2* gene is located, is significantly higher than in other non-coding regions of the *Y. hyperborea* genome (Figure S4). Moreover, it is statistically unlikely that such a sequence in the genome is so accidentally similar to the *COX2* gene (Figure 3C). All this indicates that we have detected the process of pseudogenization of one of the paralogs of the *COX2* gene in the female mitochondrial genome of *Y. hyperborea*.

**Figure 10.**
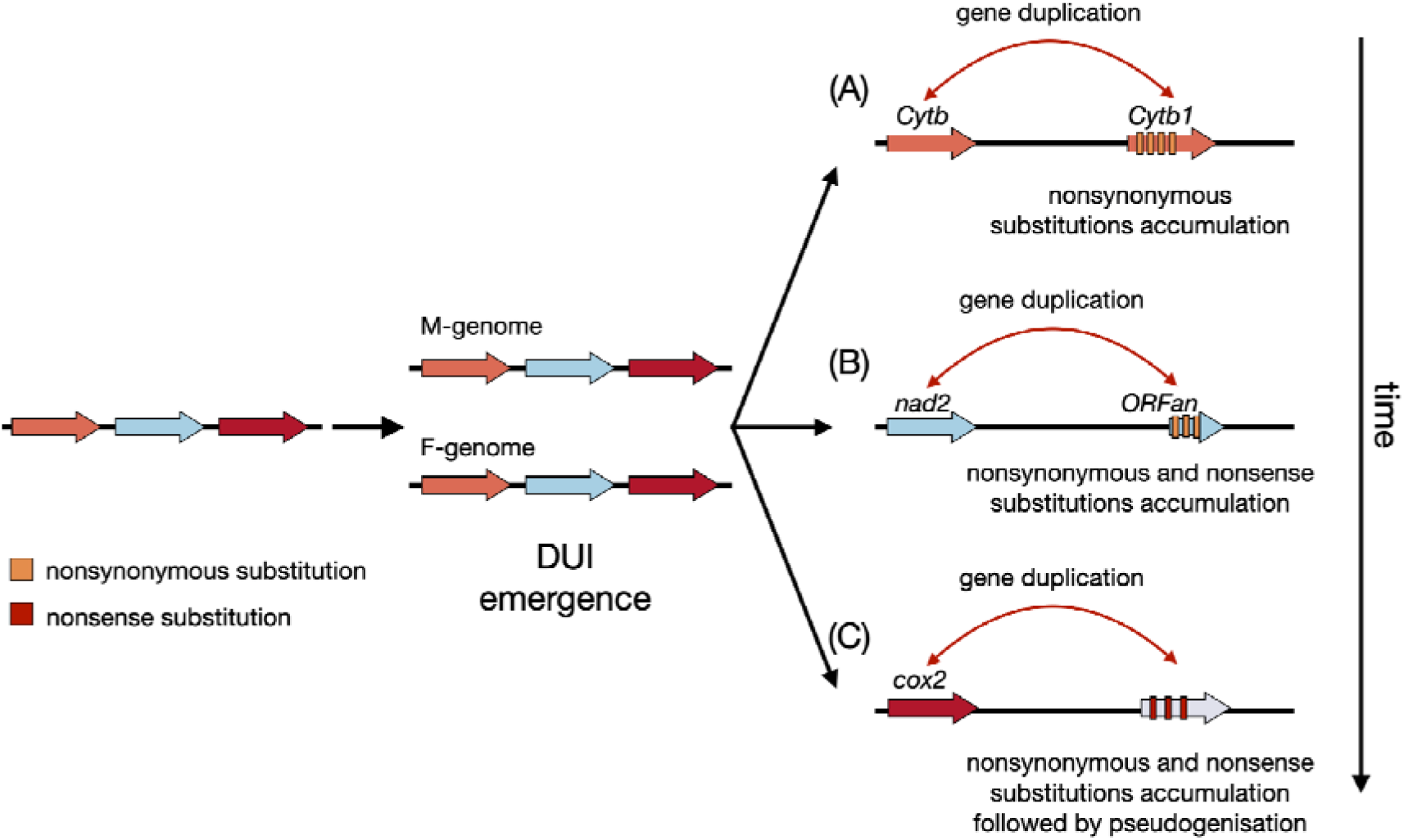
Proposed mechanism of mitochondrial genome evolution following DUI emergence. (A) Gene duplication followed by accumulation of nonsynonymous mutations, which is observed in the female mitochondrial genome of *Y. hyperborea* (*cytB* gene). (B) Gene duplication followed by accumulation of nonsense (stop) and nonsynonymous mutations, which resulted in a highly divergent ORFan gene, whose homology to *nad2* can be revealed using blast. (C) Gene duplication followed by accumulation of multiple nonsense mutations which resulted in pseudogenization of one copy of duplicated gene. This situation is observed in the female mitochondrial genome of *Y. hyperborea* (*cox2* gene).

In summary, the mitochondrial genomes of *Y. hyperborea* contain at least three regions that are degraded to varying degrees by mutational processes and relaxed selection. We cannot determine why these three genes exhibit differing levels of preservation: this variability may be linked to the timing of duplication events or to distinct evolutionary trajectories influenced by their genomic positions and functional roles. Nonetheless, our study provides a unique insight into gene duplication processes and the rapid evolution of one copy in recently emerged M- and F- type mitochondrial genomes of a basal bivalve mollusk. Rapid evolution process involves accumulation of the non-synonymous mutations, nonsense mutations, and indels in one of the duplicated gene copies.

## Materials and methods

### Specimen collection and deposition

The mollusks used in the present study were collected in the summer of 2024 in Kandalaksha Bay of the White Sea. Sampling was provided during several cruises of RV Professor Zenkevich using Van Veen (0.1 m^2^) grab from depths 60-120 m. Mature females of *Y. hyperborea* were found from the end of June to the middle of August, and mature females were found in July to August. Parts of gonads were dissected and placed into RNA- later (Evrogen) and frozen at -80° C. Each individual was preserved in 75% ethanol and deposited in the collection of the Zoological Museum of Moscow State University; part of the tissue (a fragment of mantle) of each individual was fixed in 96% ethanol for genetic analysis.

### DNA and RNA extraction, library preparation and sequencing

DNA from samples WS16905 (male) and WS16906 (female) was extracted using a Diatom DNA kit (Isogen) according to the manufacturer’s recommendations. DNA libraries were constructed using the NEBNext Ultra II DNA Library Prep Kit by New England Biolabs (NEB, MA, USA) and the NEBNext Multiplex Oligos for Illumina (Index Primers Set 1) by NEB following the manufacturer’s protocol. The samples were amplified using ten cycles of polymerase chain reaction (PCR). The constructed libraries were sequenced on an Illumina HiSeq4000 with a paired-end read length of 150.

RNA from samples WS20682 (male) and WS30680 (female) was extracted using an ExtractRNA kit (Evrogen) according to the manufacturer’s recommendations. cDNA libraries for Illumina sequencing were constructed using the NEBNext Ultra II RNA Library Prep Kit for Illumina (NEB, MA, USA) following the manufacturer’s protocol. The samples were amplified using fifteen cycles of polymerase chain reaction (PCR). cDNA libraries were sequenced on an Illumina NextSeq500 with a paired-end read length of 150.

### Assembly and analysis of mitochondrial genomes and transcriptomes

The quality of the obtained raw genomic and transcriptomic Illumina reads was analysed with fastQC v.0.12.0 followed by trimming in Trimmomatic v.0.39 (Bolger et al. 2014). Genomic ONT reads quality was analysed with NanoPlot v.1.42.0 followed by NanoFilt v. 1.42.0 trimming by length and quality (De Coster and Rademakers 2023). For the genome assembly we used a hybrid genomic mode in Spades assembler v. 3.15.5 with both Illumina and ONT reads as an input (Prjibelski et al. 2020). In order to assemble male mitochondrial genome of *Y. hyperborea* we firstly designed primers for regions which were highly diverged compared to females (See File S1). Then, using Illumina libraries obtained from long-range PCR with these primers we assembled genome with Spades assembler v. 3.15.5 using the default mode (Prjibelski et al. 2020) Mitochondrial genome annotation was performed in Mitos2 (Bernt et al. 2013). Obtained annotations were manually verified by the BLAST search (Altschul et al. 1990). Coverage was estimated by back-mapping reads to the obtained genomic sequence. Illumina reads were mapped with bowtie2 v.2.4.4 (Langmead and Salzberg 2012), while ONT reads were mapped with minimap2 v.2.23 (Li 2018). Obtained files were processed with samtools v.1.3.5 (Li et al. 2009) to obtain the exact coverage values. All obtained genomic sequences were deposited in the Genbank under the following IDs: *Yoldia hyperborea* male: PP541907; *Yoldia hyperborea* female: PQ164718; *Nuculana pernula*:(BankIt submission ID: 2966903).

Transcriptomes from RNAseq reads were assembled in rnaSPAdes v.3.15.2 (Bushmanova et al. 2019) and Trinity v.2.13.2 (Grabherr et al. 2011). Nucleotide sequences of mitochondrial proteins and mitochondrial RNAs were used as a query, while assembled transcriptome contigs were used as a database in BLAST search, using tblastx or blastn. The distribution of top blast hits was analysed and best assembly was used in further analysis. The multiple alignment of mitochondrial proteins was constructed with clustalw2 (Thompson et al. 1994). Obtained alignment was concatenated with seqkit (Shen et al. 2016). Concatenated alignment of all mitochondrial proteins except ATPase subunit 8 was further used in order to reconstruct the ML phylogenetic tree in iqtree2 v.2.3.4 (Minh et al. 2020) with a preliminary assessment of the substitution model (Kalyaanamoorthy et al. 2017). ORFans used in phylogenetic tree construction were obtained from a previous study (Breton et al. 2009).

For RNA-seq coverage estimation, Illumina reads were back-mapped to the genomic sequences using hisat2 v.2.2.1 (Kim et al. 2019). The obtained files were processed with samtools v.1.3.5 (Li et al. 2009) to obtain the exact coverage values (samtools depth option). To compare the expression level of male and female genes variants in transcriptomes, both transcriptome libraries (male and female) were separately back mapped on the previously assembled mitochondrial genomes simultaneously with hisat2 v.2.2.1 (Kim et al. 2019). For comparison, expression levels in female gonads were normalised by dividing all values by 0.9006 to account for the smaller total number of reads in this preparation.

### Identification of pseudogenized cytochrome c oxidase II in female *Yoldia* genome

We aligned two complete mitochondrial genomes of *Y. hyperborea* (male and female) using MUSCLE with default parameters (UGENE version 40.1). We then selected the aligned sequence, which corresponds to the *COX2* gene in the female genome. The obtained sequence was used for further analysis. Next, we generated 1,000 of random sequences with the same GC content as in the male *COX2* gene. Each sequence was aligned with the female region corresponding to *COX2* gene and the similarity score was counted as percent of similar positions in alignment. In addition, we have generated 1,000 sequences by codon reshuffling from the male *COX2* gene. Each reshuffled sequence was again aligned to the female region corresponding to *COX2* gene and similarity score was recorded.

### Sequence polymorphism and dN/dS calculation

For polymorphism estimation, genomes of DUI species were categorized as male or female (supplementary file 2). For each M-F pair, proteins were aligned with clustalw2 (Thompson et al. 1994). The obtained pairwise protein alignment with corresponding gene sequence was used for pal2nal v.14 alignment (Suyama et al. 2006). Using the pairwise codon alignment, the p-distance was calculated for each mitochondrial gene separately using a biopython script (Armano and Manconi 2009).

For intrapopulation variability estimation, transcriptomic reads from both male and female libraries were back mapped onto the reference male and female genomes using hisat2 v.2.2.1 (Kim et al. 2019). Since male and female transcriptomes contain both female and male variants, we could directly measure the polymorphism between the two female genomes. Four types of genomic sequences were generated using samtools v.1.3.5 (Li et al. 2009) (samtools consensus option) and mapped reads: male genes in the male transcriptome, female genes in the male transcriptome, male genes in the female transcriptome and female genes in the female transcriptome. Since male genes in the female transcriptome are poorly covered, we used only highly expressed genes for further polymorphism estimations (*COX1, COX2, COX3, rRNAL*). The obtained gene sequences were aligned using BLAST (Altschul et al. 1990) and a fraction of polymorphic positions were calculated.

For the phylogenetic tree construction, *COX1* protein was used from 53 species of bivalves (see supplementary file 2). Clustalw2 used for alignment and iqtree2 v.2.3.4 (Minh et al. 2020) with the model mtInv+F+I+G4 (Kalyaanamoorthy et al. 2017) and 1,000 ultrafast bootstrap replicates (Hoang et al. 2018) were used for ML phylogenetic tree construction. Alignment of mitochondrial proteins was performed in clustalw2 followed by pal2nal v.14 codon alignment. The obtained codon alignments were used for dN/dS estimation in codeml paml v.4.10.7 (Yang 2007). Three types of models in codeml were used: 0 model estimated mean dN/dS for each mitochondrial gene separate; model 1 (free branch model) estimated dN/dS for each branch of the tree separately; and model 2 (branch model) with selected branches were used to compare dN/dS with the model 0 estimation. Two types of phylogenetic trees with separate branches were used (see supplementary file 3): one with the *Yoldia hyperborea* M-mitogenome selected (for the *CYTB* gene test), and another with all male branches selected.

## Authors contributions

DF, GK and AT obtained and identified the specimens; ME and TN isolated DNA, prepared the libraries, performed NGS sequencing, and verified ORFans with Sanger sequencing; DF, AB and DK performed computational analysis and prepared the illustrations. DF, GB and DK drafted manuscript text; GB, TN, and DK supervised the project; TN obtained funding; All authors participated in conceptualisation, manuscript writing, and approved the final version of the manuscript.

## Data availability statement

All sequences obtained and analysed in the current study were deposited in NCBI Genbank with ID’s indicated in File S3, list “Species used ID’s” including *de novo* assembled and annotated species (PP541907.1, PQ164718.1, Submitted Bantkit: 2966903).

Conflict of Interest

## Supplementary files

**File S1.** Primers designed for male and female genomes of *Y. hyperborea*.

**File S2.** The phylogenetic tree used for synonymous and nonsynonymous substitution estimation with paml.

**File S3**. All numerical data used in the article for graphs construction and hypothesis testing with detailed description of each dataset sheet

**Figure S1.** Multiple alignment of ORFan and *NAD2* sequences of Mytilidae and Protobranchia.

**Figure S2.** Y. hyperborea RNA-seq coverage of the male genome. The outer track represents the boundaries of standard mitochondrial protein coding genes, the ORFan, tRNAs and rRNAs. Arrow direction corresponds to the coding strand.

**Figure S3.** *Y.hyperborea* RNA-seq coverage of female genome. The outer track represents the boundaries of standard mitochondrial protein coding genes, tRNAs and rRNAs. Arrow direction corresponds to the coding strand.

**Figure S4.** Mitochondrial protein divergence between male and female variants.

**Figure S5.** Scatterplot showing the relationship between divergence of F- and M-genomes and the average dN/dS (ω) in protein-coding genes of bivalve species with divergent mitochondrial genomes.

**Supplementary table S1.** Table of ORFan blast hit distribution from the male genome of Y. hyperborea against the complete genomes of other bivalves. In most cases, it is clear that the orphan finds homology in regions that correspond to the nad2 gene in different species of mollusks.

**Supplementary table S2.** The Omega values estimated for the entire phylogenetic tree (w model 0) and separately for the terminal branches corresponding to the male mitogenomes, terminal branches corresponding to the female mitogenomes, and the remaining (terminal or internal) branches. The log-likelihood values and the corresponding chi-square value is shown.

**Supplementary table S3.** The dN/dS values, estimated for all protein coding genes separately. W model 1 column shows the values for the whole branches and w model 2 columns show M-specific and rest omega values for branch-type specific codeml model. The likelihood logarithm and corresponding chi-square value is shown. In all statistical tests the number of parameters for the first model is 105 and 106 for the second, respectively.

## References

Altschul SF, Gish W, Miller W, Myers EW, Lipman DJ. 1990. Basic local alignment search tool. J. Mol. Biol. 215:403–410.

Armano G, Manconi A. 2009. ProDaMa: an open source Python library to generate protein structure datasets. BMC Res. Notes 2:202.

Bernt M, Donath A, Jühling F, Externbrink F, Florentz C, Fritzsch G, Pütz J, Middendorf M, Stadler PF. 2013. MITOS: improved de novo metazoan mitochondrial genome annotation. Mol. Phylogenet. Evol. 69:313–319.

Bettinazzi S, Nadarajah S, Dalpé A, Milani L, Blier PU, Breton S. 2020. Linking paternally inherited mtDNA variants and sperm performance. Philos. Trans. R. Soc. Lond. B Biol. Sci. 375:20190177.

Birky CW Jr. 2001. The inheritance of genes in mitochondria and chloroplasts: laws, mechanisms, and models. Annu. Rev. Genet. 35:125–148.

Bolger AM, Lohse M, Usadel B. 2014. Trimmomatic: a flexible trimmer for Illumina sequence data. Bioinformatics 30:2114–2120.

Boyle EE, Etter RJ. 2013. Heteroplasmy in a deep-sea protobranch bivalve suggests an ancient origin of doubly uniparental inheritance of mitochondria in Bivalvia. Mar. Biol. 160:413–422.

Breton S, Beaupré HD, Stewart DT, Hoeh WR, Blier PU. 2007. The unusual system of doubly uniparental inheritance of mtDNA: isn’t one enough? Trends Genet. 23:465–474.

Breton S, Beaupré HD, Stewart DT, Piontkivska H, Karmakar M, Bogan AE, Blier PU, Hoeh WR. 2009. Comparative mitochondrial genomics of freshwater mussels (Bivalvia: Unionoida) with doubly uniparental inheritance of mtDNA: gender-specific open reading frames and putative origins of replication. Genetics 183:1575–1589.

Breton S, Stewart DT, Shepardson S, Trdan RJ, Bogan AE, Chapman EG, Ruminas AJ, Piontkivska H, Hoeh WR. 2011. Novel protein genes in animal mtDNA: a new sex determination system in freshwater mussels (Bivalvia: Unionoida)? Mol. Biol. Evol. 28:1645–1659.

Burskaia V, Naumenko S, Schelkunov M, Bedulina D, Neretina T, Kondrashov A, Yampolsky L, Bazykin GA. 2020. Excessive parallelism in protein evolution of lake Baikal amphipod species flock. Genome Biol. Evol. 12:1493–1503.

Burzyński A, Śmietanka B, Fernández-Pérez J, Lubośny M. 2024. The absence of canonical respiratory complex I subunits in male-type mitogenomes of three Donax species. Sci. Rep. 14:14465.

Bushmanova E, Antipov D, Lapidus A, Prjibelski AD. 2019. rnaSPAdes: a de novo transcriptome assembler and its application to RNA-Seq data. Gigascience [Internet] 8. Available from: 10.1093/gigascience/giz100

Cao L, Kenchington E, Zouros E. 2004. Differential segregation patterns of sperm mitochondria in embryos of the blue mussel (Mytilus edulis). Genetics 166:883–894.

Capt C, Bouvet K, Guerra D, Robicheau BM, Stewart DT, Pante E, Breton S. 2020. Unorthodox features in two venerid bivalves with doubly uniparental inheritance of mitochondria. Sci. Rep. 10:1087.

Crouch NMA, Edie SM, Collins KS, Bieler R, Jablonski D. 2021. Calibrating phylogenies assuming bifurcation or budding alters inferred macroevolutionary dynamics in a densely sampled phylogeny of bivalve families. Proc. Biol. Sci. 288:20212178.

Curole JP, Kocher TD. 2002. Ancient sex-specific extension of the cytochrome c oxidase II gene in bivalves and the fidelity of doubly-uniparental inheritance. Mol. Biol. Evol. 19:1323–1328.

De Coster W, Rademakers R. 2023. NanoPack2: population-scale evaluation of long-read sequencing data. Bioinformatics [Internet] 39. Available from: 10.1093/bioinformatics/btad311

Dégletagne C, Abele D, Glöckner G, Alric B, Gruber H, Held C. 2021. Presence of male mitochondria in somatic tissues and their functional importance at the whole animal level in the marine bivalve Arctica islandica. *Commun*. Biol. 4:1104.

Dégletagne C, Abele D, Held C. 2016. A distinct mitochondrial genome with DUI-like inheritance in the ocean quahog Arctica islandica. Mol. Biol. Evol. 33:375–383.

Doucet-Beaupré H, Breton S, Chapman EG, Blier PU, Bogan AE, Stewart DT, Hoeh WR. 2010. Mitochondrial phylogenomics of the Bivalvia (Mollusca): searching for the origin and mitogenomic correlates of doubly uniparental inheritance of mtDNA. BMC Evol. Biol. 10:50.

Fisher C, Skibinski DOF. 1990. Sex-biased mitochondrial DNA heteroplasmy in the marine musselMytilus. Proc. Biol. Sci. 242:149–156.

Grabherr MG, Haas BJ, Yassour M, Levin JZ, Thompson DA, Amit I, Adiconis X, Fan L, Raychowdhury R, Zeng Q, et al. 2011. Full-length transcriptome assembly from RNA-Seq data without a reference genome. Nat. Biotechnol. 29:644–652.

Guerra D, Plazzi F, Stewart DT, Bogan AE, Hoeh WR, Breton S. 2017. Evolution of sex-dependent mtDNA transmission in freshwater mussels (Bivalvia: Unionida). Sci. Rep. 7:1551.

Gusman A, Lecomte S, Stewart DT, Passamonti M, Breton S. 2016. Pursuing the quest for better understanding the taxonomic distribution of the system of doubly uniparental inheritance of mtDNA. PeerJ 4:e2760.

Hoang DT, Chernomor O, von Haeseler A, Minh BQ, Vinh LS. 2018. UFBoot2: Improving the Ultrafast Bootstrap Approximation. Mol. Biol. Evol. 35:518–522.

Hoeh WR, Blakley KH, Brown WM. 1991. Heteroplasmy suggests limited biparental inheritance of Mytilus mitochondrial DNA. Science 251:1488–1490.

Hoeh WR, Stewart DT, Guttman SI. 2002. High fidelity of mitochondrial genome transmission under the doubly uniparental mode of inheritance in freshwater mussels (Bivalvia: Unionoidea). Evolution 56:2252–2261.

Hoeh WR, Stewart DT, Saavedra C, Sutherland BW, Zouros E. 1997. Phylogenetic evidence for role-reversals of gender-associated mitochondrial DNA in Mytilus (Bivalvia: Mytilidae). Mol. Biol. Evol. 14:959–967.

Hoeh WR, Stewart DT, Sutherland BW, Zouros E. 1996. Multiple origins of gender-associated mitochondrial DNA lineages in bivalves (Mollusca: Bivalvia). Evolution 50:2276–2286.

Kalyaanamoorthy S, Minh BQ, Wong TKF, von Haeseler A, Jermiin LS. 2017. ModelFinder: fast model selection for accurate phylogenetic estimates. Nat. Methods 14:587–589.

Kim D, Paggi JM, Park C, Bennett C, Salzberg SL. 2019. Graph-based genome alignment and genotyping with HISAT2 and HISAT-genotype. Nat. Biotechnol. 37:907–915.

Kolmogorov M, Yuan J, Lin Y, Pevzner PA. 2019. Assembly of long, error-prone reads using repeat graphs. Nat. Biotechnol. 37:540–546.

Kryazhimskiy S, Plotkin JB. 2008. The population genetics of dN/dS. PLoS Genet. 4:e1000304.

Kvist L, Martens J, Nazarenko AA, Orell M. 2003. Paternal leakage of mitochondrial DNA in the great tit (Parus major). Mol. Biol. Evol. 20:243–247.

Ladoukakis ED, Zouros E. 2001. Direct evidence for homologous recombination in mussel (Mytilus galloprovincialis) mitochondrial DNA. Mol. Biol. Evol. 18:1168–1175.

Langmead B, Salzberg SL. 2012. Fast gapped-read alignment with Bowtie 2. Nat. Methods 9:357–359.

Lee W, Zamudio-Ochoa A, Buchel G, Podlesniy P, Marti Gutierrez N, Puigròs M, Calderon A, Tang H-Y, Li L, Mikhalchenko A, et al. 2023. Molecular basis for maternal inheritance of human mitochondrial DNA. Nat. Genet. 55:1632–1639.

Li H. 2018. Minimap2: pairwise alignment for nucleotide sequences. Bioinformatics 34:3094– 3100.

Li H, Handsaker B, Wysoker A, Fennell T, Ruan J, Homer N, Marth G, Abecasis G, Durbin R, 1000 Genome Project Data Processing Subgroup. 2009. The Sequence Alignment/Map format and SAMtools. Bioinformatics 25:2078–2079.

Lubośny M, Przyłucka A, Śmietanka B, Burzyński A. 2020. Semimytilus algosus: first known hermaphroditic mussel with doubly uniparental inheritance of mitochondrial DNA. Sci. Rep. 10:11256.

Lynch M, Conery JS. 2000. The evolutionary fate and consequences of duplicate genes. Science 290:1151–1155.

Mastrantonio V, Latrofa MS, Porretta D, Lia RP, Parisi A, Iatta R, Dantas-Torres F, Otranto D, Urbanelli S. 2019. Paternal leakage and mtDNA heteroplasmy in Rhipicephalus spp. ticks. Sci. Rep. 9:1460.

Milani L, Ghiselli F, Guerra D, Breton S, Passamonti M. 2013. A comparative analysis of mitochondrial ORFans: new clues on their origin and role in species with doubly uniparental inheritance of mitochondria. Genome Biol. Evol. 5:1408–1434.

Minh BQ, Schmidt HA, Chernomor O, Schrempf D, Woodhams MD, von Haeseler A, Lanfear R. 2020. IQ-TREE 2: New Models and Efficient Methods for Phylogenetic Inference in the Genomic Era. Mol. Biol. Evol. 37:1530–1534.

Mizi A, Zouros E, Moschonas N, Rodakis GC. 2005. The complete maternal and paternal mitochondrial genomes of the Mediterranean mussel Mytilus galloprovincialis: implications for the doubly uniparental inheritance mode of mtDNA. Mol. Biol. Evol. 22:952–967.

Mugal CF, Wolf JBW, Kaj I. 2014. Why time matters: codon evolution and the temporal dynamics of dN/dS. Mol. Biol. Evol. 31:212–231.

Ort BS, Pogson GH. 2007. Molecular population genetics of the male and female mitochondrial DNA molecules of the California sea mussel, Mytilus californianus. Genetics 177:1087–1099.

Prjibelski A, Antipov D, Meleshko D, Lapidus A, Korobeynikov A. 2020. Using SPAdes De Novo Assembler. Curr. Protoc. Bioinformatics 70:e102.

Rawson PD, Hilbish TJ. 1995. Evolutionary relationships among the male and female mitochondrial DNA lineages in the Mytilus edulis species complex. Mol. Biol. Evol. 12:893–901.

Rocha EPC, Smith JM, Hurst LD, Holden MTG, Cooper JE, Smith NH, Feil EJ. 2006. Comparisons of dN/dS are time dependent for closely related bacterial genomes. J. Theor. Biol. 239:226–235.

Sano N, Obata M, Komaru A. 2007. Quantitation of the male and female types of mitochondrial DNA in a blue mussel, Mytilus galloprovincialis, using real-time polymerase chain reaction assay. Dev. Growth Differ. 49:67–72.

Sato M, Sato K. 2013. Maternal inheritance of mitochondrial DNA by diverse mechanisms to eliminate paternal mitochondrial DNA. Biochim. Biophys. Acta 1833:1979–1984.

Sato M, Sato K, Tomura K, Kosako H, Sato K. 2018. The autophagy receptor ALLO-1 and the IKKE-1 kinase control clearance of paternal mitochondria in Caenorhabditis elegans. Nat. Cell Biol. 20:81–91.

Shen W, Le S, Li Y, Hu F. 2016. SeqKit: A Cross-Platform and Ultrafast Toolkit for FASTA/Q File Manipulation. PLoS One 11:e0163962.

Skibinski DO, Gallagher C, Beynon CM. 1994. Mitochondrial DNA inheritance. Nature 368:817–818.

Stoletzki N, Eyre-Walker A. 2011. The positive correlation between dN/dS and dS in mammals is due to runs of adjacent substitutions. Mol. Biol. Evol. 28:1371–1380.

Stolyarova AV, Bazykin GA, Neretina TV, Kondrashov AS. 2019. Bursts of amino acid replacements in protein evolution. R. Soc. Open Sci. 6:181095.

Suyama M, Torrents D, Bork P. 2006. PAL2NAL: robust conversion of protein sequence alignments into the corresponding codon alignments. Nucleic Acids Res. 34:W609– W612.

Tassé M, Choquette T, Angers A, Stewart DT, Pante E, Breton S. 2022. The longest mitochondrial protein in metazoans is encoded by the male-transmitted mitogenome of the bivalve Scrobicularia plana. Biol. Lett. 18:20220122.

Thompson JD, Higgins DG, Gibson TJ. 1994. CLUSTAL W: improving the sensitivity of progressive multiple sequence alignment through sequence weighting, position-specific gap penalties and weight matrix choice. Nucleic Acids Res. 22:4673–4680.

Venetis C, Theologidis I, Zouros E, Rodakis GC. 2006. No evidence for presence of maternal mitochondrial DNA in the sperm of Mytilus galloprovincialis males. Proc. Biol. Sci. 273:2483–2489.

Walker BJ, Abeel T, Shea T, Priest M, Abouelliel A, Sakthikumar S, Cuomo CA, Zeng Q, Wortman J, Young SK, et al. 2014. Pilon: an integrated tool for comprehensive microbial variant detection and genome assembly improvement. PLoS One 9:e112963.

Wolf JBW, Künstner A, Nam K, Jakobsson M, Ellegren H. 2009. Nonlinear dynamics of nonsynonymous (dN) and synonymous (dS) substitution rates affects inference of selection. Genome Biol. Evol. 1:308–319.

Xu T, He C, Han X, Kong L, Li Q. 2024. Comparative mitogenomic analysis and phylogeny of Veneridae with doubly uniparental inheritance. Open Biol. 14:240186.

Yang Z. 2007. PAML 4: phylogenetic analysis by maximum likelihood. Mol. Biol. Evol. 24:1586–1591.

Zouros E, Oberhauser Ball A, Saavedra C, Freeman KR. 1994. An unusual type of mitochondrial DNA inheritance in the blue mussel Mytilus. Proc. Natl. Acad. Sci. U. S. A. 91:7463–7467.

