## Supplementary material for "Rapid divergence of male and female mitochondrial genomes in a basal protobranch bivalve *Yoldia hyperborea*": Figure S2.docx

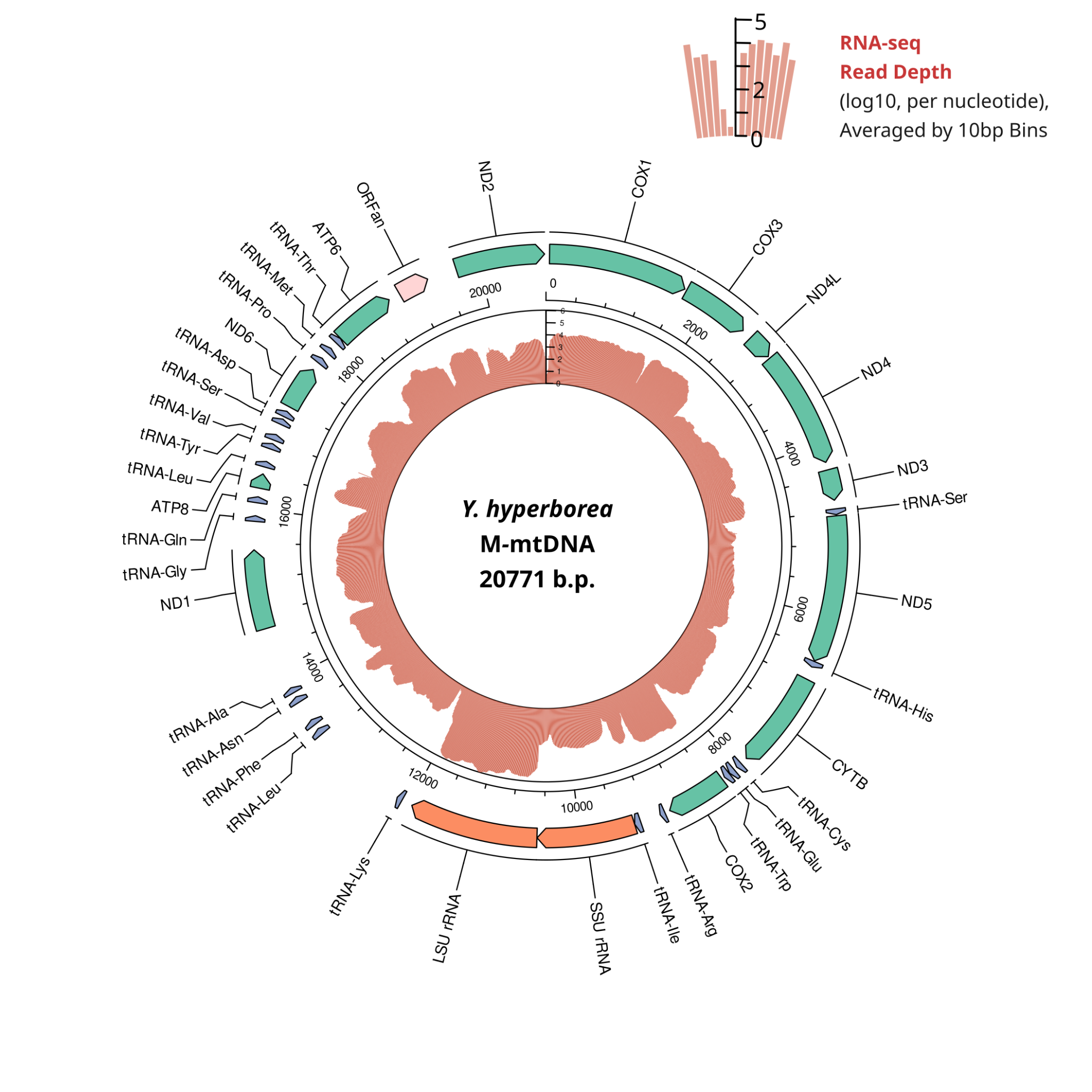

### **Figure S2.** *Y. hyperborea* RNA-seq coverage of the male genome. The outer track represents the boundaries of standard mitochondrial protein coding genes, the ORFan, tRNAs and rRNAs. Arrow direction corresponds to the coding strand.
