## Supplementary material for "Rapid divergence of male and female mitochondrial genomes in a basal protobranch bivalve *Yoldia hyperborea*": Figure S3.docx

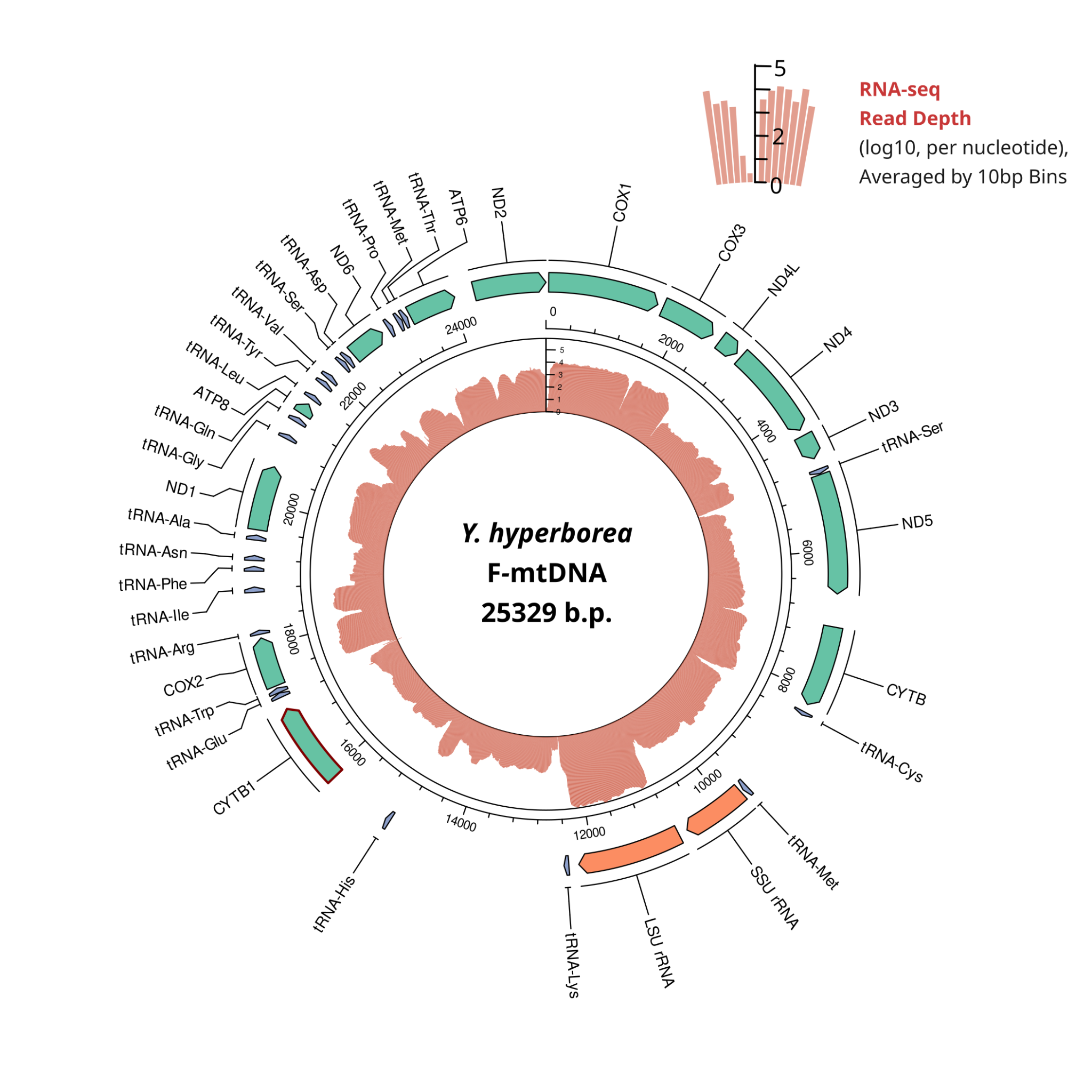

### **Figure S3.** *Y.hyperborea* RNA-seq coverage of female genome. The outer track represents the boundaries of standard mitochondrial protein coding genes, tRNAs and rRNAs. Arrow direction corresponds to the coding strand.
