## Supplementary figures and images for "Rapid divergence of male and female mitochondrial genomes in a basal protobranch bivalve *Yoldia hyperborea*"

### Figure S1.docx

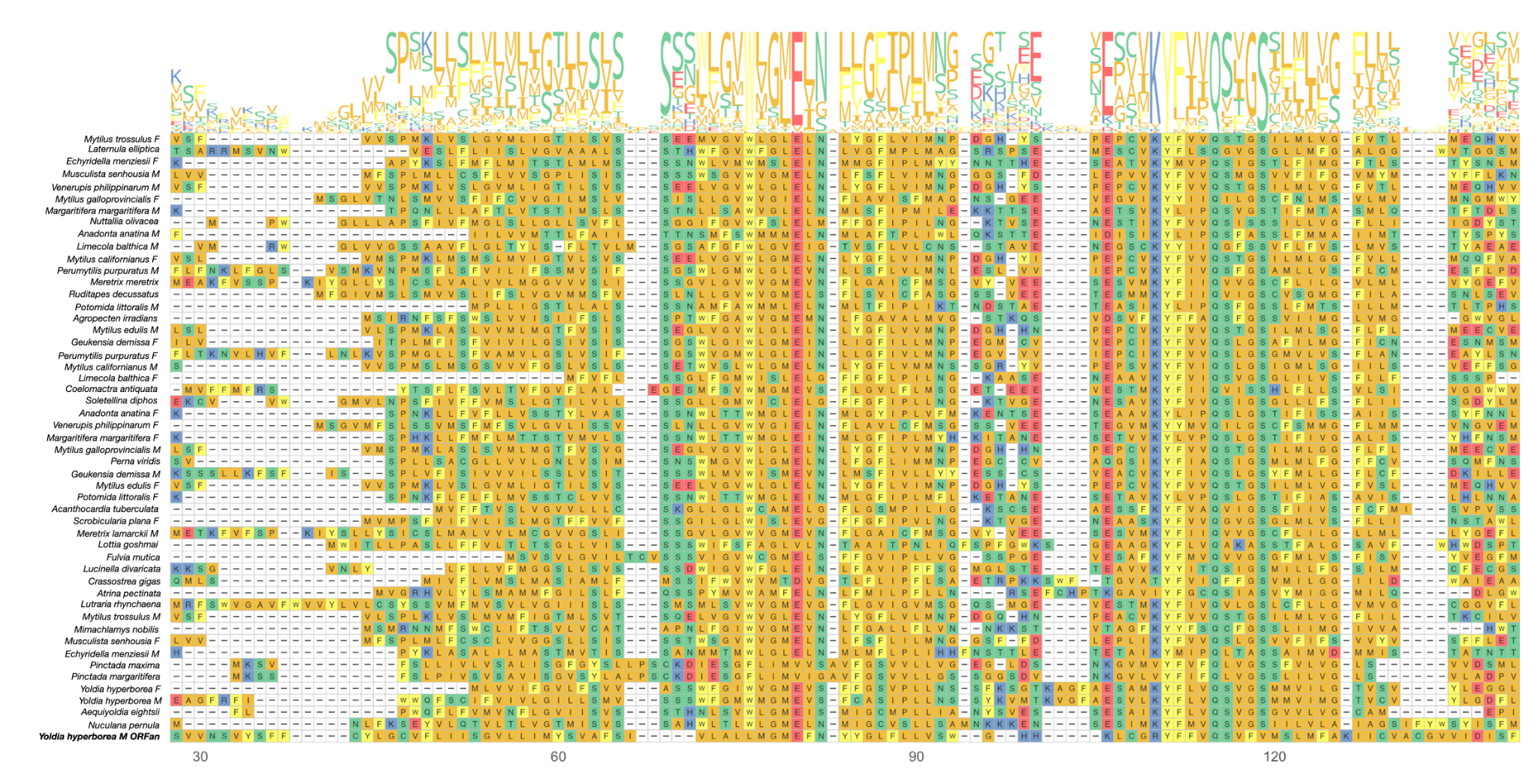

#### **Figure S1.** Multiple alignment of ORFan and *NAD2* sequences of Mytilidae and Protobranchia.

### Figure S4.docx

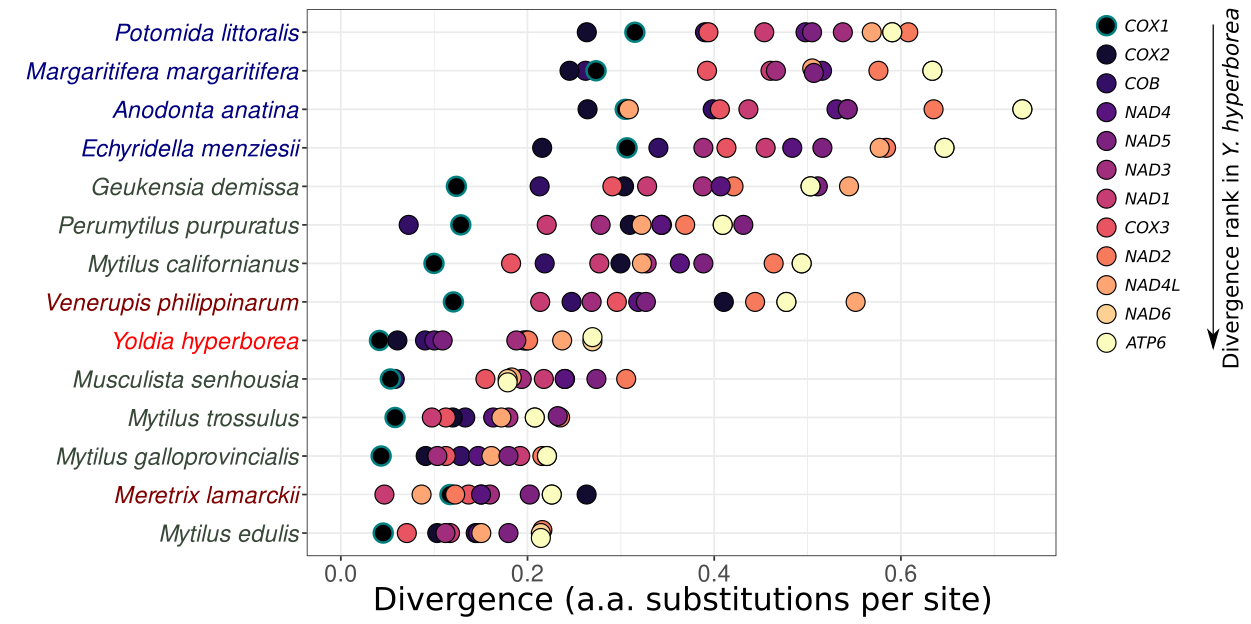

#### **Figure S4.** Mitochondrial protein divergence between male and female variants.
